# NeuroMark-SZ: A Holistic Resting-State-fMRI-Based Model for Divergent Functional Circuitry in Schizophrenia

**DOI:** 10.64898/2026.03.12.710902

**Authors:** Kyle M. Jensen, Ram Ballem, Spencer Kinsey, Pablo Andrés-Camazón, Zening Fu, Jiayu Chen, Shalaila S. Haas, Covadonga M. Díaz-Caneja, Juan R. Bustillo, Adrian Preda, Theo G.M. van Erp, Godfrey Pearlson, Peter Kochunov, Jessica A. Turner, Vince D. Calhoun, Armin Iraji

**Author notes:** Corresponding author: Kyle M. Jensen, Georgia State University, 55 Park Place, 18th Floor – TReNDS Center, Atlanta, GA 30303.

## Abstract

**Background:** Schizophrenia is a severe neuropsychiatric disorder. Efforts to describe the underlying biology and establish diagnostic markers through non-invasive neuroimaging methods are ongoing, resulting in a range of theoretical brain-based frameworks. Prominent frameworks for aberrant schizophrenia-associated functional connectivity in resting-state functional magnetic resonance imaging (rsfMRI) include the dysconnectivity hypothesis, theory of cognitive dysmetria, and triple network theory. Although informative, prior work can be improved by increasing sample size, avoiding confirmation bias, and accounting for individual variability and the effects of medication and chronicity.

**Methods:** With these recommendations in mind, we employed a data-driven, whole-brain approach using a large multi-site rsfMRI dataset (*N* = 2,656; schizophrenia = 1,248). We used reference-guided independent component analysis to generate subject-specific whole-brain functional network connectivity (FNC) and extract imaging markers of similarity to schizophrenia patterns. We modeled the relationship between medication dosage, age of onset, chronicity, symptom severity, and cognitive performance and FNC.

**Results:** Our analysis identified a reliable schizophrenia-FNC signature characterized by aberrantly stronger negative cerebellothalamic and positive thalamocortical connectivity, implicating sensory, motor, and associative cortical circuits. While medication and chronicity were significantly associated with these signatures, the core cerebellothalamic disruptions remained a robust marker of schizophrenia.

**Conclusions:** This work represents the largest schizophrenia-specific rsfMRI study to date, refines existing theoretical frameworks with a more nuanced map of how clinical variables interact with brain connectivity, and provides a high-fidelity template of schizophrenia-related connectivity. We have released this template as an open-access resource to facilitate reproducibility and accelerate the development of reliable rsfMRI-based schizophrenia biomarkers.

## 1. Introduction

Schizophrenia is a neuropsychiatric disorder with profound and far-reaching effects for both individuals and society (1). Treatment for schizophrenia can be challenging due to delayed or inaccurate diagnosis (2,3) as well as suboptimal medication efficacy and adherence (4–6). This is further compounded by the clinical heterogeneity of schizophrenia, including differences in symptoms and cognitive deficits (7,8). Resting-state functional magnetic resonance imaging (rsfMRI)(9) holds potential to improve diagnostic accuracy and identify modifiable targets by advancing our understanding of the underlying neurobiology of schizophrenia (10–12). Several theoretical frameworks have been proposed to account for key patterns observed in rsfMRI (13,14). We aim to highlight three prominent rsfMRI-based frameworks, namely the dysconnectivity hypothesis (15,16) the cognitive dysmetria or cerebello-thalamo-cortical model (17), and the triple network model (14,18), to compare them against the results of a set of statistically powerful data-driven analyses in a large multi-site dataset, and to identify their links with clinical and cognitive measures.

### 1.1. rsfMRI-Based Frameworks for Schizophrenia

The dysconnectivity hypothesis proposed by Friston (15,16) attributes the etiology of schizophrenia to disruptions in brain circuitry, leading to aberrant connectivity or “dysconnectivity”. This framework is supported by the schizophrenia-associated patterns of functional connectivity observed in rsfMRI, although its interpretations range from dysconnectivity in specific regions such as the thalamus and prefrontal cortex (19–21)(**Figure 1A**), to widespread dysconnectivity throughout the whole brain (22–24)(**Figure 1B**). The dysconnectivity hypothesis is often referenced in conjunction with the theory of cognitive dysmetria (17), implicating disruptions in cortical-subcortical-cerebellar circuitry or more specifically cerebello-thalamo-cortical connectivity (13,19) as a prototypical trait of schizophrenia. Although a large body of research provides support for these frameworks (13,19,24–33), many of the specific patterns appear to lack consistency in both directionality (i.e., positive or negative) and the regions implicated.

**Figure 1:**
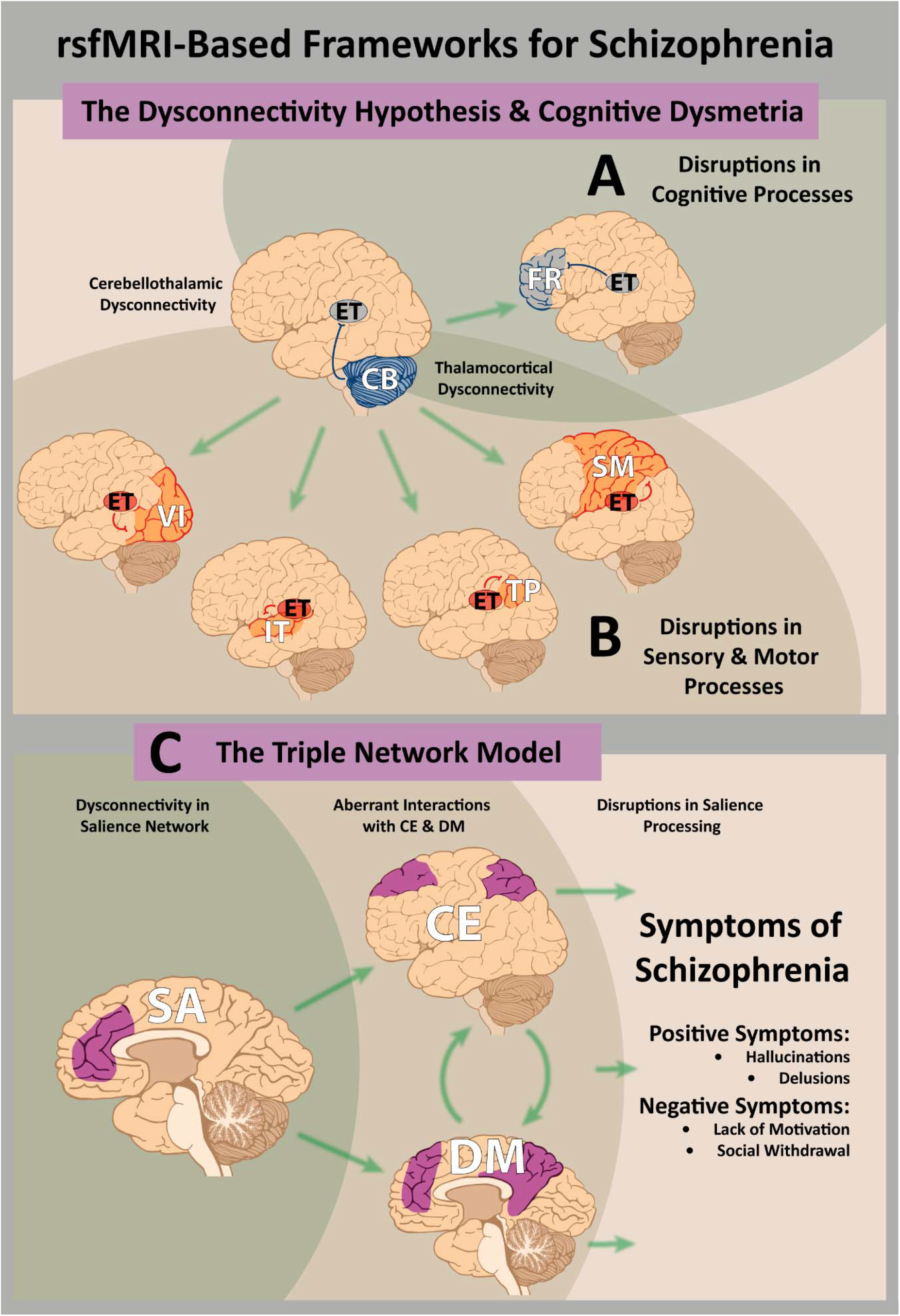
rsfMRI-Based Frameworks for Schizophrenia. The first panel illustrates how disruptions in cerebellothalamic circuitry may result in disruptions in critical modulatory processes, which in turn may lead to thalamocortical dysconnectivity, giving rise to the various symptoms of schizophrenia. Theorized efferent connections are portrayed extending from the cerebellum to the thalamus and from the thalamus to various cortical regions which spatially overlap with the NeuroMark subdomains. The NeuroMark subdomain labels are used to demonstrate how previous literature projects into the same reference space used in the current study: cerebellar (CB), visual (VI), extended thalamic (ET), sensorimotor (SM), insular-temporal (IT), temporoparietal (TP), and frontal (FR). Efferent connections portrayed in red indicate that aberrantly stronger positive connectivity is observed in schizophrenia and efferent connections portrayed in blue indicate that aberrantly stronger negative connectivity is observed in schizophrenia. **(A)** The dysconnectivity hypothesis and theory of cognitive dysmetria have traditionally focused on disruptions in prefrontal cortex and its implications for disruptions in cognitive processes. **(B)** Many recent studies report hyperconnectivity between the thalamus and sensory and motor cortex, suggesting that these patterns are relevant to observed disruptions in sensory and motor processes. The second panel **(C)** illustrates how disruptions in connectivity within the triple network domain may result in the symptoms of schizophrenia. Specifically, dysconnectivity within the salience (SA) network, comprised of the anterior cingulate cortex and anterior insula, leads to aberrant interactions with (and between) the central executive (CE) network, comprised of the dorsolateral prefrontal cortex and posterior parietal cortex, and the default mode (DM) network, comprised of the posterior cingulate cortex, precuneus, and medial prefrontal cortex, resulting in disrupted salience processing which in turn may give rise to the positive and negative symptoms of schizophrenia.

A review by Jensen and colleagues (27) suggests that although many investigators emphasize the prefrontal cortex as a critical node, the more consistently reported patterns of static functional connectivity include hypoconnectivity (defined as connectivity with negative directionality in schizophrenia relative to controls) between the cerebellum and thalamus accompanied by hyperconnectivity (positive directionality relative to controls) between the thalamus and sensorimotor, visual, and auditory cortex. Additionally, they highlight hyperconnectivity between subcortical structures in the basal ganglia and sensorimotor and temporal cortex, hypoconnectivity between cortical temporal regions, and hyperconnectivity between the cerebellum and sensorimotor and temporal cortex. Many of these patterns have been reported previously in other schizophrenia studies as well, particularly cerebellothalamic dysconnectivity (19,32,34), weaker thalamic-prefrontal connectivity (19,32,35–37), stronger thalamic-somatosensory connectivity (19,32,35,37–39), stronger thalamic-visual connectivity (32,39,40), and stronger thalamic-insular-temporal connectivity (32) in individuals with schizophrenia.

The Triple Network model is another theoretical framework that incorporates the dysconnectivity hypothesis (14,18). It is centered on three well-established large-scale resting-state networks: the frontoparietal central executive network (CEN), the default mode network (DMN), and the salience network (SN)(18)(**Figure 1C**). The CEN functions as a top-down attention control mechanism for problem-solving and decision-making (18), the DMN is associated with self-referential thought processes (18,41–43), and the SN is considered the structural substrate of bottom-up attention processing as it detects relevant – or salient – information and triggers a shift in attention, thus acting as a mediator between the externally directed CEN and internally directed DMN (18,44). The Triple Network model posits that aberrant activation, generally hypoconnectivity (45), of the SN contributes to the symptoms of schizophrenia, as a misattribution of salience might explain disorganized thought processes or the departure from reality experienced in psychosis (14). For example, if the brain incorrectly assigns significance to external stimuli or internal mental events, it may over- or under-emphasize the stimuli, resulting in maladaptive sensory processing (14). Disruptions in these three large-scale networks have been implicated as key markers of schizophrenia (45–58).

Although each framework offers a unique perspective, they are not entirely independent of each other. Furthermore, these frameworks appear to be underspecified, explaining only a fraction of the patterns of aberrant connectivity reported across studies. A whole-brain data-driven approach may provide insight into how these established frameworks fit together to form a more holistic model of schizophrenia.

### 1.2. Limitations of Prior Work

Prior studies have mostly used small sample sizes (*N* < 100), limiting the detection of smaller effect sizes (59). Additionally, while hypothesis-driven research is fundamental to science (34,39,60), it may inadvertently introduce bias and some effects can only be uncovered by considering whole-brain connectivity patterns in data-driven analyses (61). Many data-driven approaches in rsfMRI have utilized independent component analysis (ICA) to identify neuronal-related signals, often referred to as functional sources (62–64) or resting-state networks (9), however, these may vary substantially from one study to the next (65), making it difficult to compare findings across studies. Moreover, many studies utilize spatially-fixed ROIs which are not sensitive to individual differences between subjects (63,66–68). Furthermore, interpretations of these results are limited by inconsistent nomenclature (69,70). Importantly, there are several clinical factors that can affect functional connectivity, including antipsychotic medications (73–75), duration of illness (76,77), and age of onset (78–82). Finally, studies might better characterize, understand, and account for heterogeneity by considering the relationship between functional connectivity and symptom and cognitive deficit severity (83). Together these factors contribute to heterogeneity and inconsistencies reported across studies (27,71,72).

### 1.3. NeuroMark: A Data-Driven Approach Towards a Holistic Model of Schizophrenia

The NeuroMark framework incorporates predefined references with data-driven approaches, aimed towards stabilizing biomarker extraction across neuroimaging analyses (61,63,65,84,85). In rsfMRI analyses, this goal has been typically accomplished by incorporating a spatial template into a fully automated pipeline with reference-guided ICA (63,65) to estimate subject-specific functional sources referred to as intrinsic connectivity networks (ICNs). Unlike spatially-fixed nodes, template-constrained ICNs are adaptive and subject-specific (63,65). Furthermore, NeuroMark ICNs integrate structures throughout the brain and reflect multifunctionality, as a given brain region may contribute to multiple sources (86). Therefore, ICNs may more appropriately represent functional sources in the brain than commonly used predefined and fixed ROIs (63,66,87). While correlations between fixed ROIs are more widely studied and typically referred to as functional connectivity, correlations between the time courses of ICNs are referred to as functional network connectivity (FNC)(67), to distinguish the ability of ICA to decompose fMRI data into both within- and between-network connectivity relative to standard ROI-based approaches (88).

The NeuroMark approach has been utilized to identify imaging markers associated with various neurological and psychiatric disorders (23,65,89–97) and multiple NeuroMark templates have been released to date (65,85,98,99). The NeuroMark 2.2 template (63,86) holds particular promise for unveiling new insights into schizophrenia, as it consists of 105 highly replicable ICNs derived from over 100,000 rsfMRI scans, encompasses the whole brain, incorporates information from across multiple spatial scales, and has demonstrated reliability across the lifespan (63,86,100). The multi-scale attribute is especially noteworthy as it enables studies to investigate relationships both within and between different spatial scales, capturing insights that might be missed by a single model order (101,102). Furthermore, NeuroMark 2.2 exhibits improved harmonization and standardization by describing functional units in terms familiar to the fields of cognitive and affective neuroscience. For example, NeuroMark 2.2 incorporates the triple network domain (86) corresponding to the large-scale networks described in Menon’s (18) triple network model. Together, these attributes make the NeuroMark approach well-equipped for addressing the aims of the current study, which seeks examine patterns of divergent whole-brain FNC in individuals with schizophrenia compared to controls in light of prominent theoretical frameworks while providing: 1) unprecedented scale enabling reliable effect size estimation, 2) enhanced spatial precision through large-sample ICA decomposition, 3) systematic quantification of medication, duration of illness, age of onset, and symptom effects in a single cohort, and 4) demonstration of robustness across clinical heterogeneity.

## 2. Methods

### 2.1. Participants

We pooled rsfMRI, demographic, clinical, and cognitive assessments from six large-scale neuroimaging datasets (**Supplement S1**) to investigate schizophrenia-control group differences in whole-brain FNC and their associations with antipsychotic use, age of onset, duration of illness, symptom severity, and cognitive performance (**Table 1**). We interpreted the results with three pre-existing theoretical frameworks and assessed their compatibility. Antipsychotic use was quantified by average daily chlorpromazine equivalence (CPZ) dosage in milligrams per day (103,104). Symptom severity was measured with the positive and negative syndrome scale (PANSS)(105) and Brief Psychiatric Rating Scale (BPRS)(106); scores of the latter were converted to PANSS based on (107). To assess cognitive performance, dataset specific cognitive assessments were harmonized into the seven cognitive domains of the Measurement and Treatment Research to Improve Cognition in Schizophrenia (MATRICS)(108) Consensus Cognitive Battery (MCCB)(109). Specific cognitive assessments for each dataset and our approach to harmonization are described in **Supplement S2**. The full sample included 1,248 individuals with schizophrenia and 1,408 controls. As reported in **Table 1**, the sex and race distributions differed significantly between the schizophrenia and control groups, but age did not. Compared to controls, the schizophrenia group also had lower cognitive performance across all cognitive domains. We statistically accounted for each of these group differences in our analyses. Subsamples of this large multi-site dataset were used in subsequent analyses, including only individuals with available data for the required variables in each.

**Table 1:**
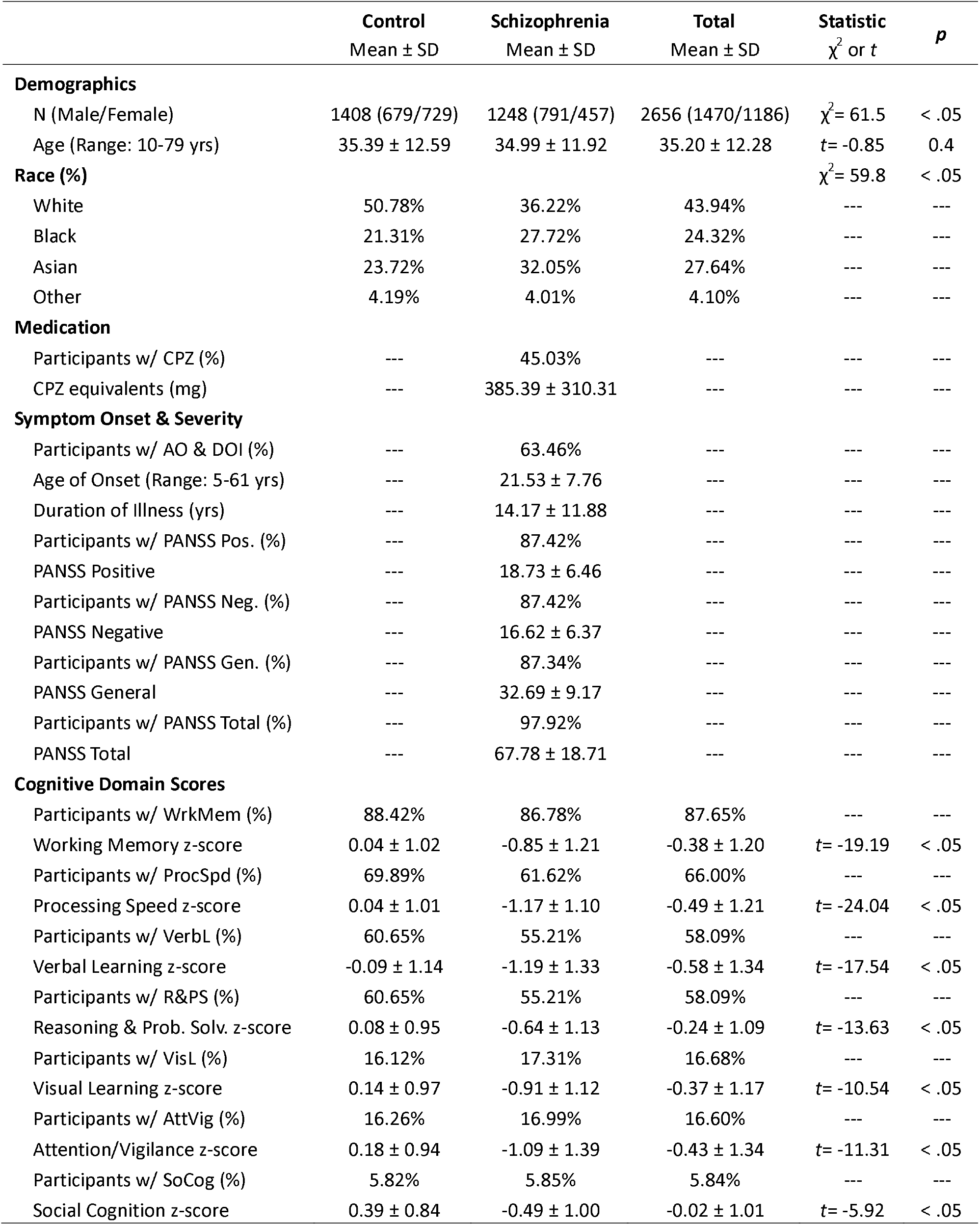
Dataset Summary. Demographic, clinical characteristics, and cognitive assessments for the combined sample are displayed. Participants are assigned to either the schizophrenia or control group. The table displays the number of participants in each group, the range, mean, and standard deviation (SD) of age in years (yrs), and the race distribution. The “Other” category for race includes participants who indicated that they belonged to a race other than those presented, indicated that they belonged to more than one race, or chose not to disclose this information. Inferential test statistics and p-values are provided where appropriate, where the control group served as the reference group (i.e., positive test statistics indicate higher values in schizophrenia relative to controls). The following clinical characteristics are reported for the schizophrenia sample: the percentage of participants with medication dosage information available, the mean chlorpromazine (CPZ) equivalent estimate in milligrams (mg) per day, the percentage of participants with age of onset (AO) and duration of illness (DOI) information available, the age of onset and duration of illness, percentage of participants with PANSS Positive (Pos.)/Negative (Neg.)/General (Gen.)/Total scores available, and the mean and SD for each. The following cognitive assessment scores are reported for all participants: the percentage of participants in each group (and total) with Working Memory (WrkMem), Processing Speed (ProcSpd), Verbal Learning (VerbL), Reasoning and Problem Solving (R&PS; Reasoning & Prob. Solv.), Visual Learning (VisL), Attention/Vigilance (AttVig), and Social Cognition (SoCog) domain scores, as well as the mean and SD of the z-scores for each test. Similar tables for each individual dataset are included in **Supplement S1**.

### 2.2. Image Acquisition and Preprocessing

Although the imaging protocols differed in spatial and temporal resolution across datasets, we utilized standardized NeuroMark preprocessing steps (63,65,85) to facilitate the comparisons made in our analyses. RsfMRI data were preprocessed in MATLAB (SPM12, https://www.fil.ion.ucl.ac.uk/spm/), including steps for discarding initial volumes, slice timing correction, a rigid body motion correction, warping to MNI space using the echo-planar imaging (EPI) template, resampling to 3×3×3 mm^3^ voxel space, spatial smoothing by a Gaussian kernel with a full width half maximum (FWHM) of 6 mm, and the variance intensity of the time courses was normalized.

### 2.3. Quality Control Validation

In each analysis, statistical tests were first performed on the full combined dataset, maximizing sample size by excluding only the participants who lacked data for the required variables. Subsequent analyses were performed in subsets with additional quality control (QC) criteria (e.g., removing participants with high head motion) consistent with prior work (63,65)(**Supplement S3** and **S4**).

### 2.4. Analyses

#### 2.4.1. Estimating Subject-Specific ICNs

To estimate subject-specific ICNs, we implemented multivariate-objective optimization ICA with reference (MOO-ICAR)(63,110) using the NeuroMark 2.2 template (86)(**Figure 2**) via the Group ICA of fMRI Toolbox (GIFT)(111).

**Figure 2:**
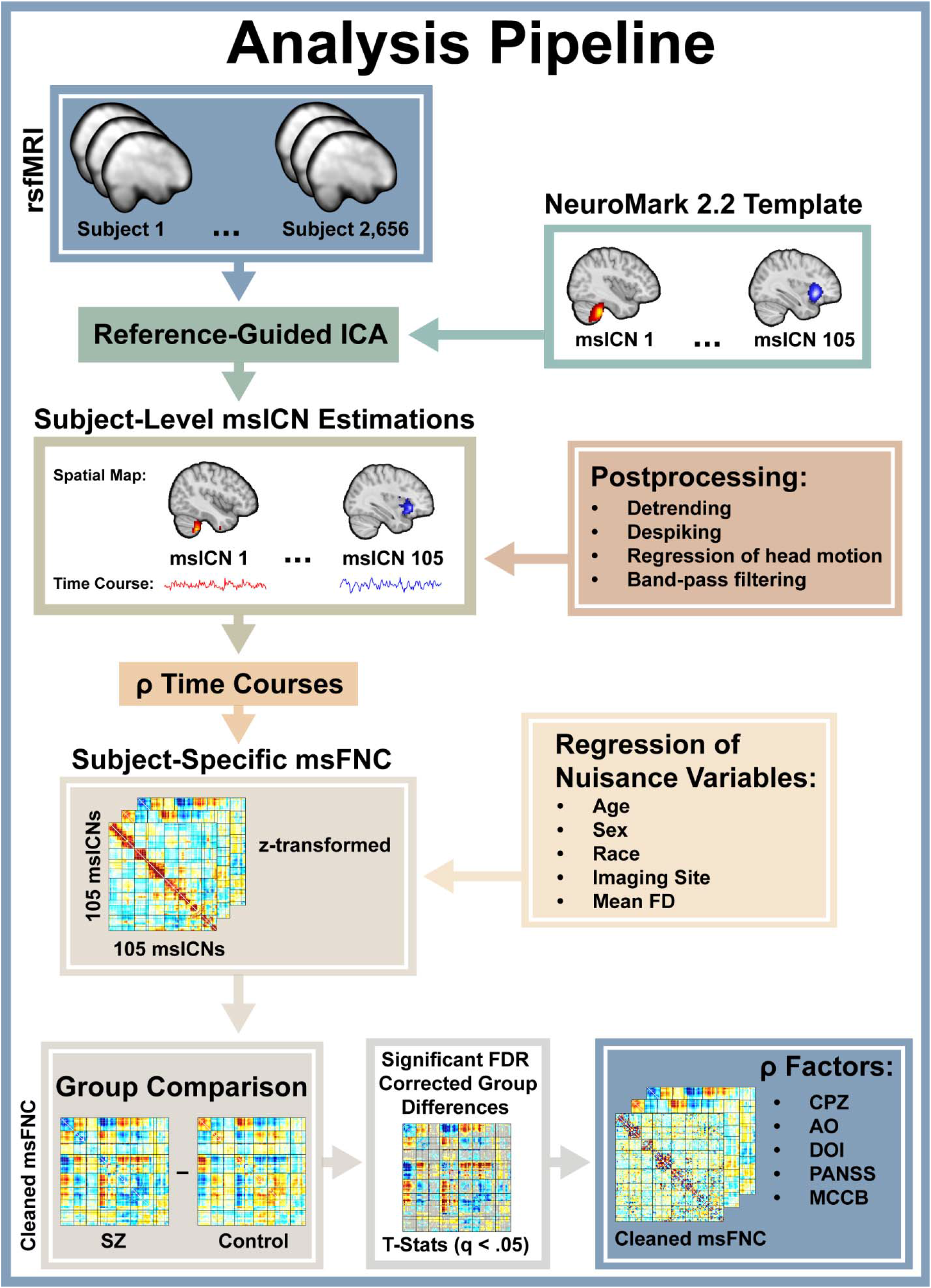
Analysis Pipeline. Reference-guided independent component analysis (ICA) is applied to preprocessed resting-state functional magnetic resonance imaging (rsfMRI) scans for 2,656 subjects with reference to the NeuroMark 2.2 multi-scale template which consists of 105 multi-scale intrinsic connectivity networks (msICNs). Individual subject-level msICNs are estimated from the rsfMRI, spatially-constrained to the NeuroMark template. Postprocessing is performed on the time courses of these 105 subject-level msICNs and then a Pearson correlation is calculated between each of them, resulting in a 105 by 105 multi-scale functional network connectivity (msFNC) matrix for each subject. Prior to statistical analyses, the msFNC is z-transformed. A cleaned msFNC is calculated by regressing out nuisance variables such as mean framewise displacement (Mean FD) and then a two-sample t-test is performed between schizophrenia (SZ) and control groups to calculate group differences. A false discovery rate (FDR) correction is performed and the msFNC pairs with significant group differences (*q* < .05) are further investigated, testing for associations with average daily chlorpromazine dose (CPZ), age of onset (AO), duration of illness (DOI), symptom severity (PANSS equivalent), and cognitive performance via MATRICS Consensus Cognitive Battery (MCCB) equivalent scores.

#### 2.4.2. Subject-Specific FNC and Group Comparisons

Postprocessing was performed on the ICN time courses consistent with previous work (65), including steps to remove additional noise effects through detrending, despiking, regression of head motion (6 motion parameters and their 6 derivatives), and 0.01-0.15 Hz band-pass filtering. We computed subject-specific FNC through a Pearson correlation between all pairs of the 105 multi-scale ICNs. Each FNC feature was z-transformed before entering a general linear model (GLM) with FNC as the response variable and the following confounding variables as covariate predictors: age, sex, race, imaging site, and mean framewise displacement (FD)(112). The raw residual variance in FNC from this GLM (i.e., cleaned FNC) were utilized in subsequent analyses. In the main analysis, a two-sample *t*-test was conducted on all features of the cleaned FNC between the schizophrenia and control groups. In the group comparison, we computed Hedge’s *g* effect sizes and adjusted for multiple comparisons using false discovery rate (FDR)(113) correction. In addition, the primary analysis was repeated for each individual imaging site, demonstrating consistency across sites and suggesting that potential site-related effects were negligible (**Supplement S5**).

#### 2.4.3. Clinical and Cognitive Associations

Using only the FNC features with significant group differences after FDR correction, we investigated the association (Pearson correlation) between the cleaned FNC and CPZ in a subset of individuals with schizophrenia who had CPZ data available. The same analysis was repeated to calculate associations with age of onset, duration of illness, and symptom subscales. A similar analysis was also performed with cognitive scores, although diagnostic group means were first subtracted from both the FNC and the cognitive scores for each subject.

## 3. Results

### 3.1. Subject-Specific ICNs and FNC

105 multi-scale ICNs were estimated for each subject and grouped into domains in accordance with NeuroMark 2.2 (**Figure 3A-N**). The average spatial maps had a mean ICN-template similarity of *r* = 0.86 (std 0.04, min 0.74, max 0.95), demonstrating high correspondence with the NeuroMark 2.2 template, while accounting for individual differences. Subject-specific FNC was extracted and averaged across subjects, displaying patterns of modularity consistent with prior studies utilizing the NeuroMark 2.2 organization (**Figure 3O**)(86,100,114).

**Figure 3:**
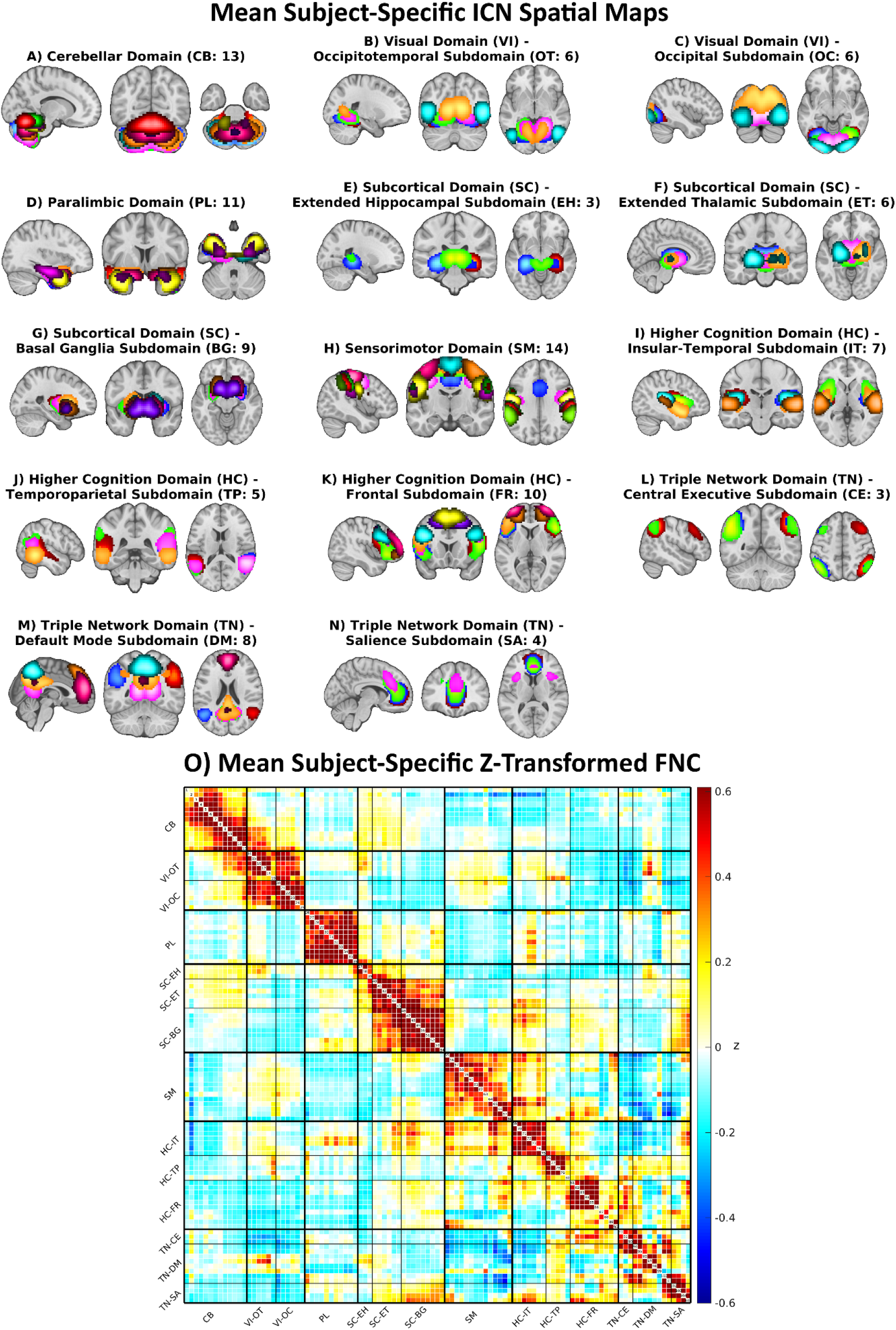
Mean ICN Spatial Maps and FNC. A composite view of the average spatial maps across all participants (*N* = 2,656) is displayed for each of the estimated 105 Intrinsic connectivity networks (ICNs). ICNs are grouped by their domain-subdomain labels, consistent with the NeuroMark_fMRI_2.2 template: **(A)** cerebellar (CB), **(B)** visual-occipitotemporal (VI-OT), **(C)** visual-occipital (VI-OC), **(D)** paralimbic (PL), **(E)** subcortical-extended hippocampal (SC-EH), **(F)** subcortical-extended thalamic (SC-ET), **(G)** subcortical-basal ganglia (SC-BG), **(H)** sensorimotor (SM), **(I)** higher cognition-insular-temporal (HC-IT), **(J)** higher cognition-temporoparietal (HC-TP), **(K)** higher cognition-frontal (HC-FR), **(L)** triple network-central executive (TN-CE), **(M)** triple network-default mode (TN-DM), and **(N)** triple network-salience (TN-SA). The spatial maps are z-scored, with voxels > 3 displayed. **(O)** Mean z-transformed functional network connectivity (FNC) for all participants is displayed in a matrix with red indicating higher positive values and blue indicating lower negative values.

### 3.2. Schizophrenia-Control Group Comparisons

Widespread disruptions in FNC were observed across the whole brain in individuals with schizophrenia (**Figure 4**). The strongest effects were negative connectivity in schizophrenia relative to controls between cerebellar and subcortical domains, specifically the extended thalamic and basal ganglia subdomains. Also notable was the stronger negative connectivity between the sensorimotor domain and the temporoparietal subdomain. These patterns were accompanied by stronger positive thalamocortical connectivity in sensory cortex (i.e., occipitotemporal, paralimbic, sensorimotor, insular-temporal, & temporoparietal). A strong pattern of positive connectivity was also observed between the cerebellar and sensorimotor domains in schizophrenia compared to controls. Notably, although many schizophrenia-control group differences survived FDR correction in the triple network domain, the effect sizes were relatively weak (mean *g* = .147 ± .048), in comparison with the domains already described (mean *g* = .239 ± .109). Moreover, the frontal subdomain displayed mixed patterns of both strong positive and negative connectivity with the thalamic subdomain in schizophrenia.

**Figure 4:**
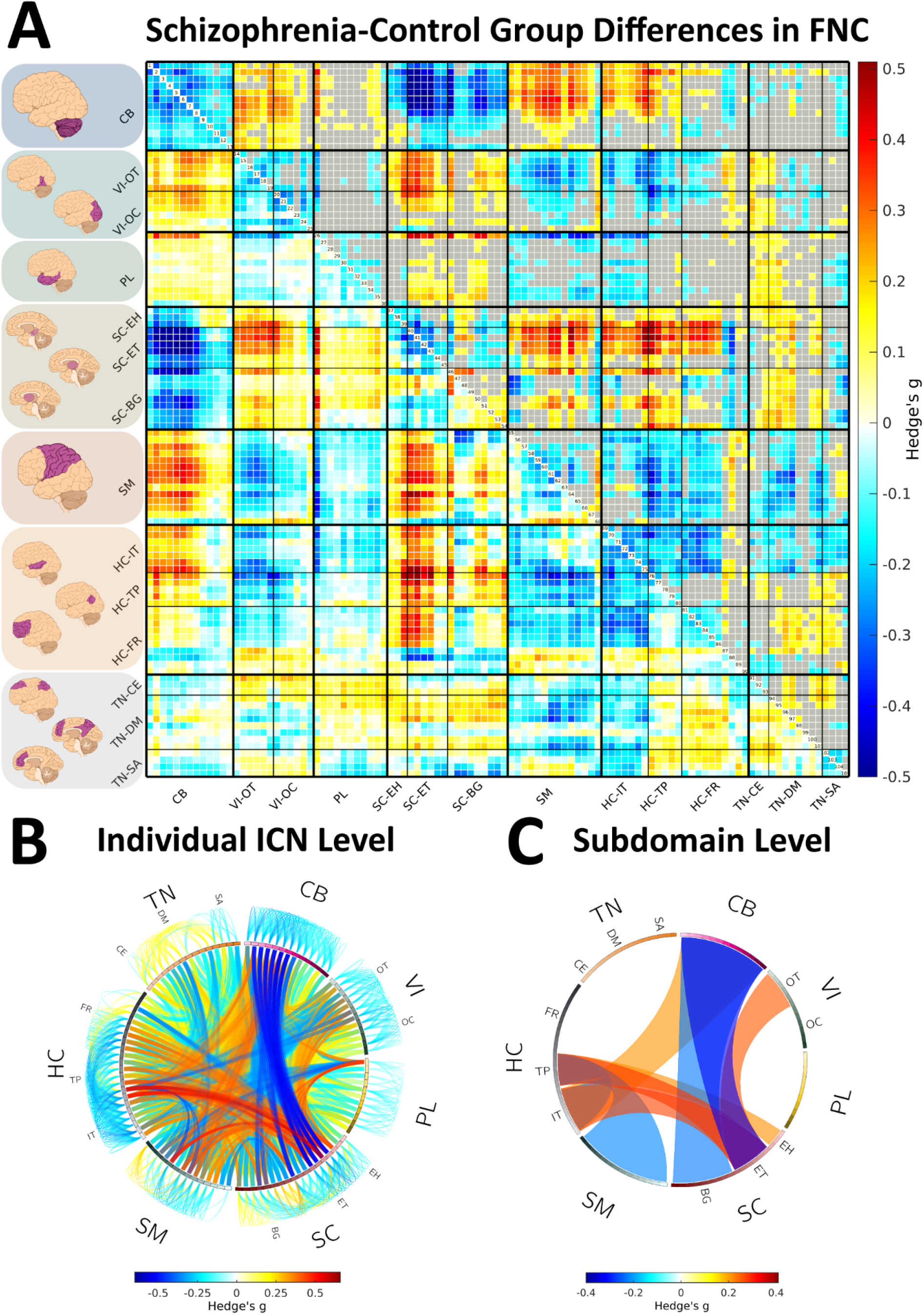
Schizophrenia-Control FNC Group Differences. Group comparisons for each combination of the 105 NeuroMark intrinsic connectivity networks (ICNs) is displayed in the matrix **(A)**. The upper triangle displays only significant (FDR corrected *q* < .05) case-control group differences in functional network connectivity (FNC) with non-significant pairs grayed out. The heatmap uses red to represent higher positive FNC and blue to represent lower negative FNC in schizophrenia compared to controls. ICNs are grouped by their domain-subdomain labels, consistent with the NeuroMark_fMRI_2.2 template: cerebellar (CB), visual-occipitotemporal (VI-OT), visual-occipital (VI-OC), paralimbic (PL), subcortical-extended hippocampal (SC-EH), subcortical-extended thalamic (SC-ET), subcortical-basal ganglia (SC-BG), sensorimotor (SM), higher cognition-insular-temporal (HC-IT), higher cognition-temporoparietal (HC-TP), higher cognition-frontal (HC-FR), triple network-central executive (TN-CE), triple network-default mode (TN-DM), and triple network-salience (TN-SA). Significant (FDR *q* < .05) patterns of FNC are also displayed in connectograms at the individual **(B)** and subdomain **(C)** levels. The subdomain level patterns represent a high-level summary by displaying the average of the significant (FDR *q* < .05) group difference effect sizes (Hedge’s *g*) for all FNC pairs within each subdomain block (e.g., OT-EH FNC), with the threshold showing only the top 10% of the strongest effects.

### 3.3. FNC and Antipsychotic Medication

Numerous associations were found between patterns of disrupted FNC in schizophrenia and CPZ dose (**Figure 5** and **Supplement S6**). Among the strongest effects were negative correlations with cerebellothalamic connectivity, accompanied by positive correlations with thalamocortical (thalamic-sensorimotor) connectivity and cerebellar-sensorimotor connectivity. To further investigate the influence of CPZ on diagnostic effect in FNC, we performed a GLM adjusting for CPZ as a covariate and found that statistically controlling for CPZ had minimal impact on case-control group differences (**Supplement S7**).

**Figure 5:**
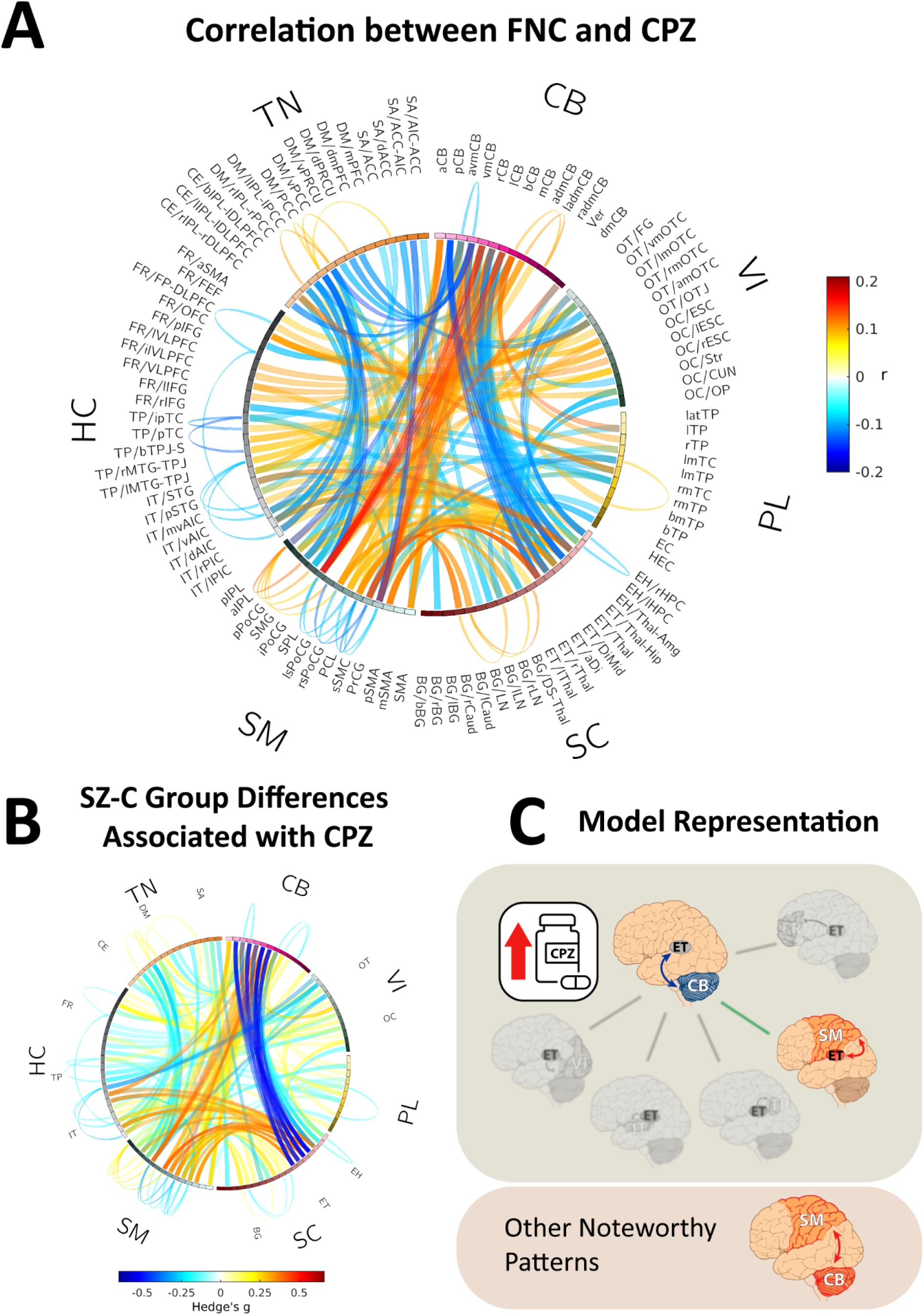
FNC and CPZ. The first connectogram **(A)** displays the correlation between functional network connectivity (FNC) and average daily chlorpromazine equivalence dosage (CPZ) for each of the FNC pairs that showed significant (FDR corrected *q* < .05) differences between the schizophrenia and control groups. Correlations were calculated for the subset of individuals with CPZ scores (*N* = 562) and only significant (*p* < .05) FNC and CPZ correlations are displayed. In this connectogram, red represents a *positive correlation*, and blue represents a *negative correlation*. ICNs are grouped by their domain-subdomain labels, consistent with the NeuroMark_fMRI_2.2 template: cerebellar (CB), visual-occipitotemporal (VI-OT), visual-occipital (VI-OC), paralimbic (PL), subcortical-extended hippocampal (SC-EH), subcortical-extended thalamic (SC-ET), subcortical-basal ganglia (SC-BG), sensorimotor (SM), higher cognition-insular-temporal (HC-IT), higher cognition-temporoparietal (HC-TP), higher cognition-frontal (HC-FR), triple network-central executive (TN-CE), triple network-default mode (TN-DM), and triple network-salience (TN-SA). The second connectogram **(B)** displays effect sizes for the significant (FDR *q* < .05) schizophrenia-control group differences in FNC from Figure 4 which had significant (*p* < .05) associations with CPZ. In this connectogram, red represents higher positive FNC and blue represents lower negative FNC in schizophrenia compared to controls. The third panel **(C)** is a visual representation of how the most prominent patterns in the current results fit into the theoretical frameworks for schizophrenia presented in Figure 1, where higher CPZ were associated with aberrant negative cerebellothalamic FNC and aberrant positive thalamic-sensorimotor FNC. Other thalamocortical patterns of FNC from the model are grayed out, as they were largely unassociated with CPZ. In addition, although previous models may not emphasize it, we also observed associations between higher CPZ and stronger positive cerebello-sensorimotor FNC.

### 3.4. FNC and Illness Onset and Duration

Age of onset was associated with FNC throughout the brain. Most notably, an older age of onset was positively correlated with thalamocortical (thalamic-paralimbic) connectivity, as well as cerebello-subcortical (basal ganglia) connectivity (**Figure 6A & 6B**). In contrast, age of onset was negatively correlated with connectivity between the salience subdomain and the insular-temporal and basal ganglia subdomains, as well as connectivity between the insular-temporal and basal ganglia subdomains. Many associations were identified between aberrant FNC in schizophrenia and duration of illness (**Figure 6C & 6D**). Among the strongest effects were negative correlations between duration of illness and thalamocortical (thalamic-temporoparietal) connectivity and visual-sensorimotor connectivity. In contrast, there was a positive correlation between duration of illness and thalamocortical (thalamic-sensorimotor & visual) connectivity.

**Figure 6:**
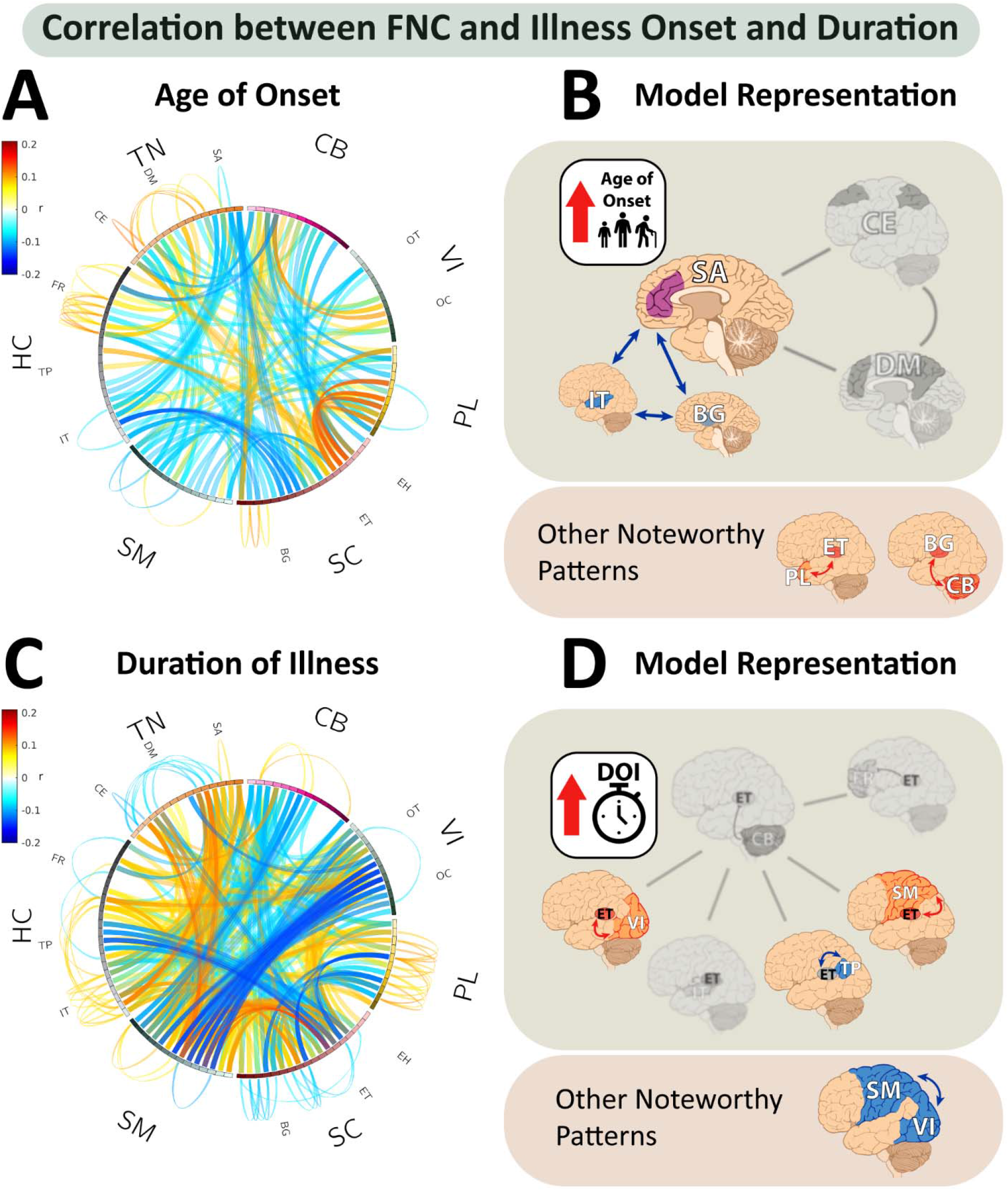
FNC and Illness Onset and Duration. The first panel **(A)** displays a connectogram of correlations between functional network connectivity (FNC) and age of onset (AO) for each of the FNC pairs with significant (FDR corrected *q* < .05) group differences between schizophrenia and control groups. The second panel **(B)** displays a visual representation of how the most prominent patterns in the current results fit into the theoretical frameworks for schizophrenia presented in Figure 1, where later AO was associated with aberrant negative salience-insular-temporal, salience-basal ganglia, and insular-temporal-basal ganglia FNC. Although previous models may not emphasize it, we also observed associations between later AO and stronger positive thalamocortical (paralimbic) and stronger negative cerebellar-basal ganglia FNC. The third panel **(C)** displays a connectogram of correlations between FNC and duration of illness (DOI) for each of the FNC pairs with significant (FDR corrected *q* < .05) group differences between schizophrenia and control groups. The fourth panel **(D)** displays a visual representation of how the most prominent patterns in the current results fit into the theoretical frameworks for schizophrenia presented in Figure 1, where a longer DOI was associated with aberrantly stronger positive thalamocortical (sensorimotor & visual) FNC and weaker positive thalamocortical (temporoparietal) FNC. Although previous models may not emphasize it, we also observed associations between longer DOI and stronger negative sensorimotor-visual FNC. In both connectograms, correlations were calculated for a subset of individuals with AO and DOI scores (*N* = 792) and only significant (*p* < .05) FNC and AO correlations are displayed. In the connectograms, red represents a *positive correlation*, and blue represents a *negative correlation*. ICNs are grouped by their domain-subdomain labels, consistent with the NeuroMark_fMRI_2.2 template: cerebellar (CB), visual-occipitotemporal (VI-OT), visual-occipital (VI-OC), paralimbic (PL), subcortical-extended hippocampal (SC-EH), subcortical-extended thalamic (SC-ET), subcortical-basal ganglia (SC-BG), sensorimotor (SM), higher cognition-insular-temporal (HC-IT), higher cognition-temporoparietal (HC-TP), higher cognition-frontal (HC-FR), triple network-central executive (TN-CE), triple network-default mode (TN-DM), and triple network-salience (TN-SA).

### 3.5. FNC and Symptom Severity

Many of the FNC pairs that showed group differences in schizophrenia were associated with symptom severity (see **Figure 7** for PANSS subscales, and **Supplement S8** for total scores). Among the strongest FNC-PANSS associations were negative correlations between thalamocortical (thalamic-sensorimotor) FNC and PANSS positive, and between thalamocortical (thalamic-visual) FNC and PANSS negative and general subscales. These were contrasted by strong positive correlations between salience-frontal and default mode-visual FNC and PANSS positive and general, between default mode-paralimbic FNC and PANSS negative, and between default mode-frontal FNC and PANSS general subscales. We also observed notable negative correlations between sensorimotor-frontal FNC and PANSS negative, and between visual-basal ganglia and cerebellocortical (cerebellar-insular-temporal) FNC and PANSS general subscales. Finally, we found a positive correlation between occipitotemporal-sensorimotor FNC and the PANSS negative subscale.

**Figure 7:**
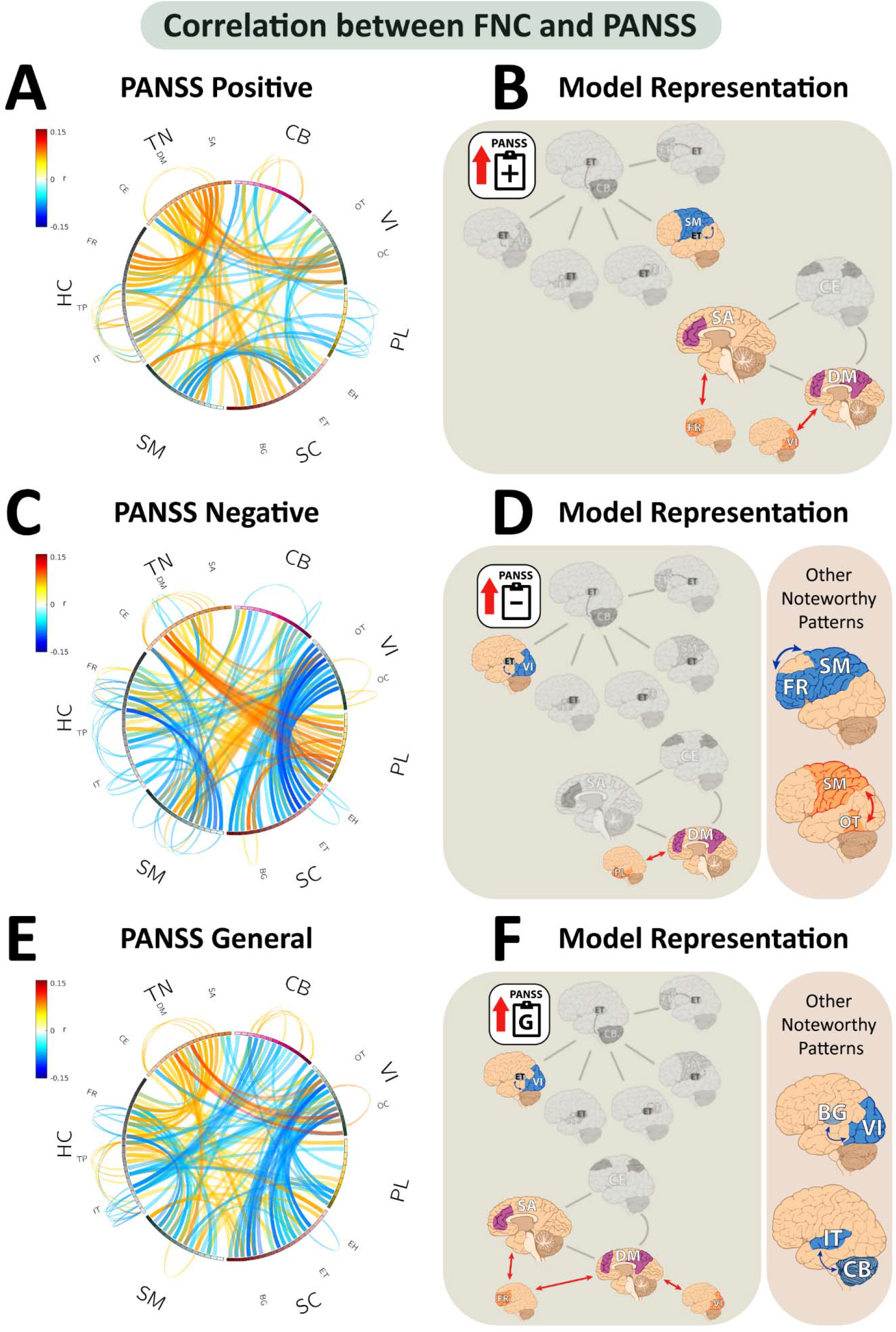
**FNC and Symptom Severity**. Each of the functional network connectivity (FNC) pairs with significant (FDR corrected *q* < .05) group differences between schizophrenia and control groups in the main analysis were tested for correlations with the PANSS positive **(A-B)**, negative **(C-D)**, and general **(E-F)** subscales, with PANSS total reported in **Figure S8**. The connectograms in the panels on the left **(A, C, E)** display significant (*p* < .05) FNC-PANSS correlations calculated for subsets of individuals with PANSS scores. In the connectograms, red represents a *positive correlation*, and blue represents a *negative correlation*. ICNs are grouped by their domain-subdomain labels, consistent with the NeuroMark_fMRI_2.2 template: cerebellar (CB), visual-occipitotemporal (VI-OT), visual-occipital (VI-OC), paralimbic (PL), subcortical-extended hippocampal (SC-EH), subcortical-extended thalamic (SC-ET), subcortical-basal ganglia (SC-BG), sensorimotor (SM), higher cognition-insular-temporal (HC-IT), higher cognition-temporoparietal (HC-TP), higher cognition-frontal (HC-FR), triple network-central executive (TN-CE), triple network-default mode (TN-DM), and triple network-salience (TN-SA). The panels on the right **(B, D, F)** visually represent how the most prominent patterns of FNC-PANSS associations relate to the theoretical frameworks for schizophrenia presented in Figure 1.

### 3.6. FNC and Cognitive Performance

Many of the FNC pairs that showed schizophrenia-control group differences were associated with cognitive performance (**Figure 8**). Although there is some overlap in patterns of associated FNC across cognitive domains, there appear to be unique functional profiles associated with each MCCB domain. Within the cognitive dysmetria and dysconnectivity hypothesis frameworks, cerebellothalamic connectivity was positively correlated with attention/vigilance, verbal learning, and visual learning. Thalamocortical (thalamic-visual) connectivity was negatively correlated with verbal learning, while thalamocortical (thalamic-insular-temporal & sensorimotor) connectivity was negatively correlated with visual learning. With regard to the triple network framework, salience-frontal connectivity was negatively correlated with attention/vigilance, default mode-frontal connectivity was negatively correlated with attention/vigilance and visual learning, default mode-basal ganglia connectivity was negatively correlated with processing speed and working memory, and default mode-sensorimotor connectivity was negatively correlated with reasoning and problem solving. Central executive-sensorimotor connectivity was also negatively correlated with reasoning and problem solving. Central executive-visual connectivity was negatively correlated with social cognition, verbal learning, and visual learning, and central executive-paralimbic connectivity was positively correlated with reasoning and problem solving and negatively correlated with visual learning. Several additional FNC associations with cognitive performance were observed that extend beyond these frameworks (**Figure 8I**).

**Figure 8:**
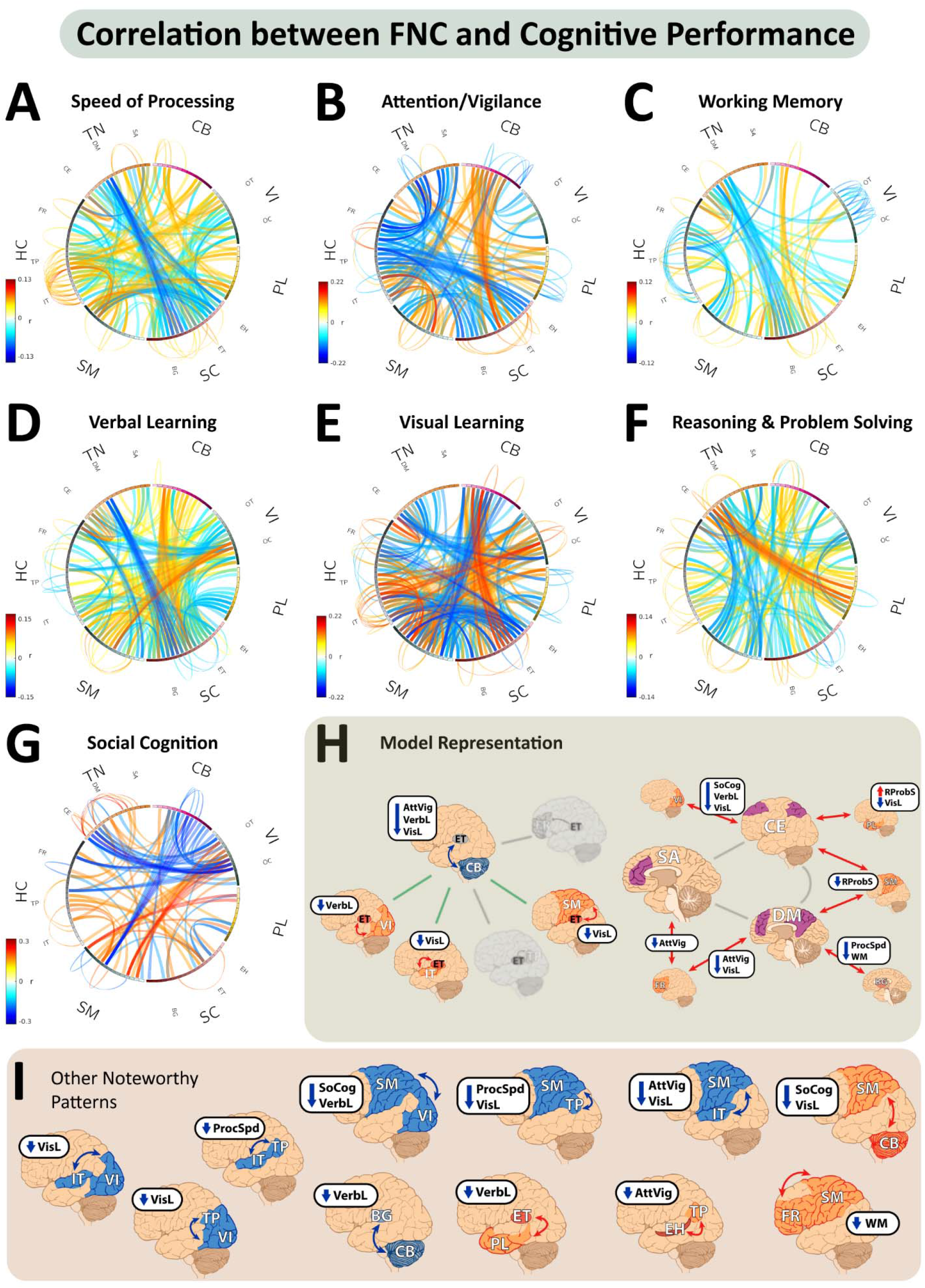
FNC and Cognitive Performance. Each of the functional network connectivity (FNC) pairs with significant (FDR corrected *q* < .05) group differences between schizophrenia and control groups in the main analysis were tested for correlations with cognitive assessment scores corresponding with seven cognitive domains. The connectograms in panels **(A-G)** display significant (*p* < .05) FNC-cognitive score correlations calculated for subsets of individuals with each score. In the connectograms, red represents a *positive correlation*, and blue represents a *negative correlation*. ICNs are grouped by their domain-subdomain labels, consistent with the NeuroMark_fMRI_2.2 template: cerebellar (CB), visual-occipitotemporal (VI-OT), visual-occipital (VI-OC), paralimbic (PL), subcortical-extended hippocampal (SC-EH), subcortical-extended thalamic (SC-ET), subcortical-basal ganglia (SC-BG), sensorimotor (SM), higher cognition-insular-temporal (HC-IT), higher cognition-temporoparietal (HC-TP), higher cognition-frontal (HC-FR), triple network-central executive (TN-CE), triple network-default mode (TN-DM), and triple network-salience (TN-SA). Panel **(H)** visually represent how the most prominent patterns of FNC-cognitive performance associations relate to the theoretical frameworks for schizophrenia presented in Figure 1. Panel **(I)** highlights some prominent FNC-cognitive performance associations which potentially extend upon existing frameworks. ProcSpd: Speed of Processing; AttVig: Attention/Vigilance; WM: Working Memory; VerbL: Verbal Learning; VisL: Visual Learning; RProbS: Reasoning & Problem Solving; SoCog: Social Cognition

### 3.7. QC Validation Analyses

We repeated the group comparison analysis in a QC subset consisting of 65% of the original sample and found a strong correlation (*r* = .97) between the effect sizes from each (**Supplement S4**). In appendage analyses, the QC validation appeared impacted by the smaller sample sizes, therefore, a post-hoc analysis randomly sampling 1,000 subsets of equal size was performed, revealing highly similar results whether QC criteria were considered or not (**Supplement S9**).

## 4. Discussion

### 4.1. Robust and Reliable Imaging Markers of Schizophrenia

Our results identified schizophrenia-control group differences in FNC throughout the whole brain with high statistical confidence. Many of the strongest effects were consistent with patterns described previously (13,15,17,19,23,27), although our results offer several new insights (**Figure 9**). Specifically, the dysconnectivity hypothesis and theory of cognitive dysmetria emphasize disruptions in the prefrontal cortex as a critical feature of schizophrenia, while disruptions in sensorimotor, paralimbic, and associative cortex are less emphasized (15–17). Similarly, triple network theory focuses on three large-scale networks (i.e., CEN, DMN, & SN) which primarily implicate frontal cortex (14,18). While these networks appear to be relevant, our results suggest that they may capture only a portion of the disrupted network activity in schizophrenia, but not the most prominent and reliable patterns. Instead, our results highlight pronounced negative cerebellar-subcortical (basal ganglia, hippocampal, thalamic) connectivity coupled with primarily positive cerebellocortical connectivity, positive subcortical-cortical connectivity, and negative cortico-cortical connectivity. Among these, aberrant thalamocortical (thalamic-paralimbic, sensorimotor, temporoparietal, insular-temporal, & occipitotemporal) connectivity was especially strong. Prior research suggests that the cerebellum modulates both motor and cognitive function through cerebellothalamic pathways (115) and disrupted connectivity in these pathways has been suggested as the key dysfunction underlying cognitive dysmetria (34). This consistently reported pattern of anticorrelated cerebellothalamic connectivity (27) may reflect a breakdown of critical modulatory processes, serving as a catalyst for aberrantly stronger positive thalamocortical connectivity affecting primary sensory, motor, and associative cortex potentially underlying the behavioral and cognitive manifestations of schizophrenia.

**Figure 9:**
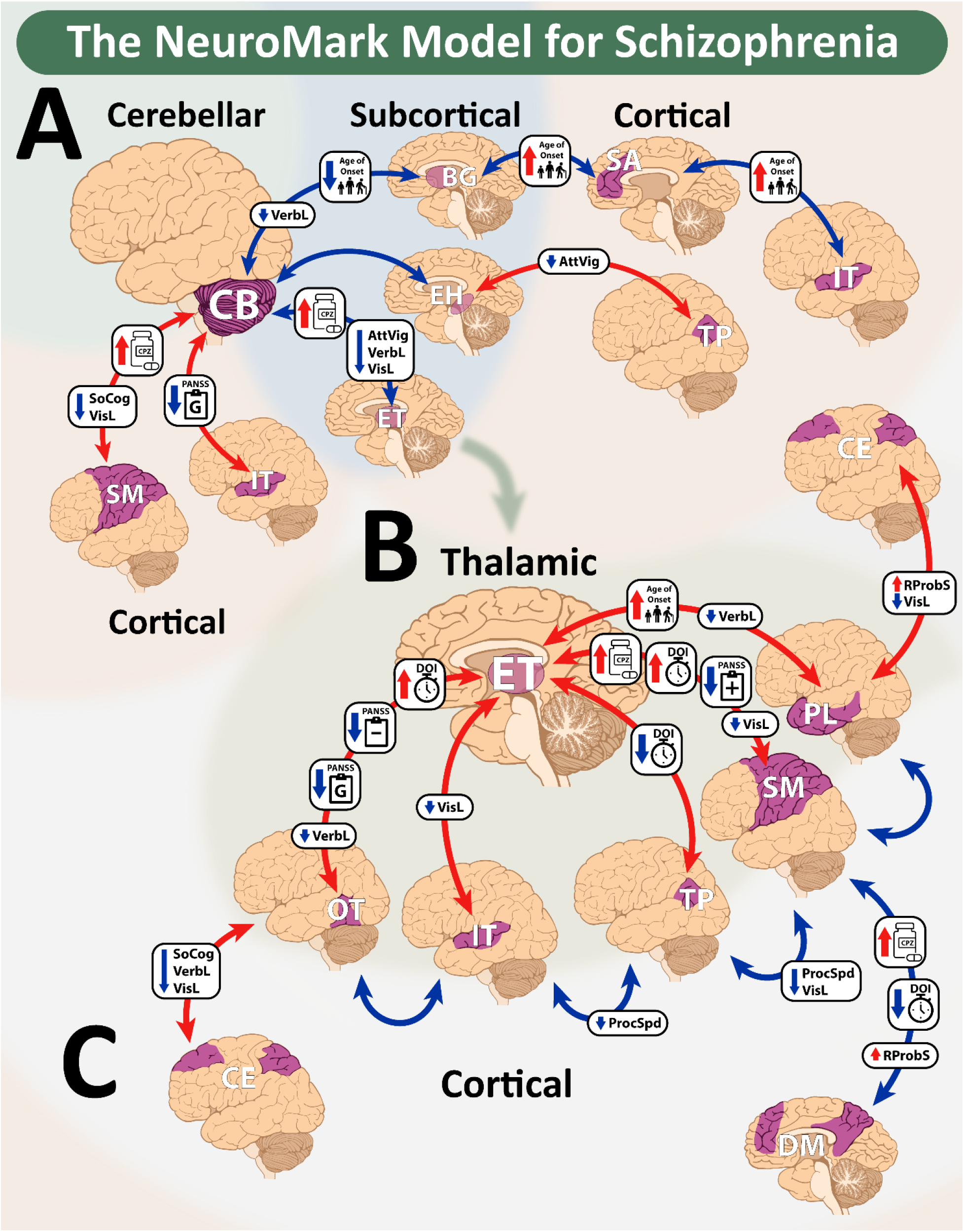
The NeuroMark Model for Schizophrenia. A collective summary of the most prominent patterns of disrupted connectivity observed in schizophrenia is shown, highlighting aberrant **(A)** cerebellar-subcortical-cortical connectivity, **(B)** thalamocortical connectivity, and **(C)** cortico-cortical connectivity. Red arrows represent *stronger positive* connectivity and blue arrows represent *stronger negative* connectivity between brain regions in the schizophrenia group relative to the control group. Associations with chlorpromazine equivalent dosage (CPZ), age of onset, duration of illness (DOI), symptom severity (PANSS), and cognitive scores are displayed along the arrows for each of these functional connectivity network (FNC) pairs. The arrows beside each variable indicate its directionality relative to the directionality of the FNC (e.g., a red arrow next to CPZ paired with a blue FNC arrow indicates that higher CPZ dosage correlates with more negative FNC). The green arrow between panels A and B serves to draw attention to the thalamus as a central node affected by schizophrenia. Each region is grouped by their domain-subdomain labels, consistent with the NeuroMark_fMRI_2.2 template: cerebellar (CB), visual-occipitotemporal (VI-OT), visual-occipital (VI-OC), paralimbic (PL), subcortical-extended hippocampal (SC-EH), subcortical-extended thalamic (SC-ET), subcortical-basal ganglia (SC-BG), sensorimotor (SM), higher cognition-insular-temporal (HC-IT), higher cognition-temporoparietal (HC-TP), higher cognition-frontal (HC-FR), triple network-central executive (TN-CE), triple network-default mode (TN-DM), and triple network-salience (TN-SA). ProcSpd: Speed of Processing; AttVig: Attention/Vigilance; WM: Working Memory; VerbL: Verbal Learning; VisL: Visual Learning; RProbS: Reasoning & Problem Solving; SoCog: Social Cognition

### 4.2. Relationships with Cognitive and Clinical Profiles

Attention/vigilance, verbal learning, and visual learning displayed strong positive correlations with cerebellothalamic dysconnectivity, suggesting that this circuitry serves as a common neural substrate for many cognitive processes, and underscoring its potential relevance to the established neuropsychological deficits in schizophrenia (36,116). Importantly, while there was some overlap in FNC-cognitive performance associations, the general trend of cognitive impairment may map onto more specific functional profiles associated with each cognitive domain, suggesting that different patterns of aberrant FNC are relevant to different cognitive deficits. For example, impaired processing speed was associated with stronger negative cortico-cortical connectivity between temporoparietal, insular-temporal, and sensorimotor cortex, regions which encompass the language-associated angular gyrus or Brodmann Area (BA) 39, and Wernicke’s Area (BA 22)(86,117).

While impaired cognitive performance generally aligned with the directionality of divergent FNC in schizophrenia, symptom severity typically displayed opposite patterns of directionality, for example, in thalamic-sensorimotor FNC. Sensorimotor-related symptoms are commonly reported in individuals with schizophrenia (118,119) and disruptions in this network-level circuitry are also frequently reported (17,120,121). Notably, much of the aberrant FNC in the thalamic-sensorimotor domains were positively related to antipsychotic dosage and duration of illness, but inversely related to positive symptoms, which may reflect the effects of antipsychotic treatments, which are generally thought to interact with dopaminergic pathways and are targeted towards treating positive symptoms (122). Nevertheless, while extrapyramidal symptoms are frequently attributed to the effects of antipsychotic medication blocking dopamine D2 receptors (122,123), alterations in sensorimotor circuitry and neurological soft signs appear to be independently related to schizophrenia as they have been observed early in the course of the disease and even prior to taking medication (124–126). Consistent with this, not all FNC pairs in the thalamic-sensorimotor domains displayed associations with antipsychotic dosage and duration of illness.

### 4.3. Limitations and Future Directions

Our findings highlight many strong effects relevant to existing theoretical frameworks of schizophrenia. To facilitate future efforts, we have released a publicly available template (**Supplement S10**) as a resource in future neuroimaging analyses aimed towards developing stable imaging markers of schizophrenia, including the creation of metrics such as individualized vulnerability indexes (e.g., see 98,129), as well as towards the development of more personalized treatment, such as cognitive training interventions (128).

Medication effects were included through translating antipsychotic doses in chlorpromazine equivalents; however, our data did not enable quantification of the effects of the different antipsychotics’ mechanisms of actions, exposure duration, and treatment response. Other potential associations and confounders that could be explored in future studies include substance use, psychotropic polypharmacy, medical comorbidity, disability severity, and other social determinants of health. The current study sought to maximize statistical power by pooling data from multiple cohorts. However, not all measures were shared across all sites, limiting some of the analyses and challenging an inherent assumption of homogeneity.

We sought to achieve a balance between maintaining scientific rigor while not excluding meaningful trends in the data due to arbitrary thresholding. Our comparisons between analyses in the full sample and QC subsets demonstrated how the magnitude of the statistical measures scaled with sample size, resulting in fewer FNC features passing the significance threshold in the QC subsets, while the patterns remained nearly identical in the primary group difference analysis and highly similar in the appendage analyses (e.g., FNC and CPZ), where sample sizes were greatly diminished. This finding suggests that our QC criteria may have been overly stringent, greatly reducing sample size and statistical power while the results were highly similar. Future studies should consider the impact of thresholding on their statistical tests and QC criteria imposed, as effect sizes are inherently small in brain-wide association studies (59). Although we demonstrated the replicability of our results in subsets of the same dataset, future investigations are needed to demonstrate the replicability of these results in independent datasets, particularly in first-episode and at-risk samples which are less confounded by or even unexposed to medication and the effects of chronicity.

Psychotic symptoms observed in schizophrenia also occur in other psychiatric disorders (7), therefore, investigating whether the current findings are also observed in other conditions with overlapping symptoms will be important regarding specificity. Additionally, subgroups with unique biological alterations may exist within any given schizophrenia cohort (89,90). Future work might consider how some of these patterns in FNC may reflect unique subgroups with distinct clinical profiles.

Lastly, there are substantial limitations regarding inferences about the neurobiological processes underlying schizophrenia based on rsfMRI findings (129). Further investigations utilizing task-based fMRI, neuromodulation, or animal models of schizophrenia could cross validate the neurobiological correlates of rsfMRI-based markers of schizophrenia (130,131).

### 4.4. Conclusions

To date, rsfMRI studies highlighted the relevance of the frontal cortex to schizophrenia through the dysconnectivity hypothesis, cognitive dysmetria, and triple network frameworks. Our results suggest that these frameworks overemphasize the robustness of aberrant connectivity in the frontal cortex as a reliable rsfMRI-based marker for schizophrenia. Instead, our results emphasize stronger negative cerebellothalamic connectivity and stronger positive thalamocortical connectivity, particularly between the thalamus and temporal cortex (occipitotemporal, insular-temporal, temporoparietal, & paralimbic), unassociated with confounds such as medication and duration of illness, and thus potentially more specific for schizophrenia. Though, further validation of these findings in first-episode, and ideally unmedicated samples is warranted. Furthermore, considering the strong associations of medication and duration of illness with specific patterns, such as stronger positive thalamic-sensorimotor connectivity, could serve as important considerations, both for study design and interpreting results. The creation and release of NeuroMark-SZ marks the beginning of an open-source data sharing initiative to construct a database of disorder-specific profiles organized within the NeuroMark framework. We anticipate that this resource will facilitate reproducibility and serve as an important aid in the process of biomarker development.

## Ethics Statement

All data were from previous studies which were conducted in accordance with the Declaration of Helsinki, were approved by their respective institutional review boards, and obtained informed consent from all participants.

## Data Availability

The NeuroMark templates described in this report are freely available at https://trendscenter.org/data/. The Group ICA of fMRI Toolbox (GIFT) is freely available at http://trendscenter.org/software/gift. The FBIRN data used in this report is publicly available (see https://doi.org/10.1016/j.neuroimage.2015.09.003). The COBRE data used in this report is publicly available from http://fcon_1000.projects.nitrc.org/indi/retro/cobre.html. Additional data which are not publicly available can be requested from the authors. Please for data and codes.

## Author Contributions

Kyle M. Jensen contributed through conceptualization, data curation, formal analysis, methodology, project administration, software, visualization, writing original draft, and writing - review & editing. Ram Ballem contributed through data curation, software, visualization, and writing - review & editing. Spencer Kinsey contributed through writing - review & editing. Pablo Andrés-Camazón contributed through conceptualization, methodology, and writing - review & editing. Zening Fu contributed through methodology, software, data curation, writing - review & editing. Jiayu Chen contributed through conceptualization, methodology, and writing - review & editing. Shalaila S. Haas contributed through conceptualization, methodology, and writing - review & editing. Covadonga M. Díaz-Caneja contributed through writing - review & editing. Juan R. Bustillo contributed through writing - review & editing. Adrian Preda contributed through writing - review & editing. Theo G.M. van Erp contributed through data curation and writing - review & editing. Godfrey Pearlson contributed through data curation and writing - review & editing. Peter Kochunov contributed through data curation and writing - review & editing. Jessica A. Turner contributed through data curation and writing - review & editing. Vince D. Calhoun contributed through conceptualization, funding acquisition, project administration, resources, supervision, and writing – review & editing. Armin Iraji contributed through conceptualization, methodology, funding acquisition, project administration, resources, supervision, and writing – review & editing.

## Acknowledgements

We thank Jing Sui (Beijing Normal University) for providing Dataset 6 (see **Supplement S1**).

