## Supplementary Information for "NeuroMark-SZ: A Holistic Resting-State-fMRI-Based Model for Divergent Functional Circuitry in Schizophrenia"

Supplementary Information S1. Dataset Descriptions

The multi-site dataset utilized in the present study includes rsfMRI scans collected from 27 different imaging sites from the FBIRN, COBRE, MPRC, B-SNIP 1, B-SNIP 2, and Dataset 6 cohorts. Descriptions of each of the six datasets including imaging protocol, recruitment details, inclusion criteria, and demographic summary tables are presented below:

FBIRN**:** The Functional Imaging Biomedical Informatics Research Network (FBIRN; Keator et al., 2016) dataset is comprised of rsfMRI collected from eight sites, each with approval from their respective Institutional Review Board (IRB). All sites utilized scanners with a 3T field strength as well as the following acquisition parameters: a standard gradient echo-planar imaging sequence, repetition and echo times (TR/TE) of 2000/30 ms, a voxel size of 3.4375 × 3.4375 × 4 mm with a 1 mm slice gap, a field of view equal to 220 × 220 mm, and a total of 162 volumes. Participants were recruited from the University of California Irvine, the University of California Los Angeles, the University of New Mexico, the University of Iowa, the University of Minnesota, Duke University/University of North Carolina, the University of California San Diego, and the University of California San Francisco. Individuals with schizophrenia completed the Structured Clinical Interview for DSM-IV-TR Axis I Disorders (First et al., 2002a, 2002b) for diagnostic confirmation and were excluded if they had a current or past history of a major medical illness, had significant extrapyramidal symptoms or tardive dyskinesia, or were not clinically stable. Participants in the control group were excluded if they had a current or past history of a major neurological or psychiatric medical illness or had a first degree relative diagnosed with a psychotic illness. All participants gave written informed consent and were between the ages of 18-62 years old, had normal hearing levels, normal vision, an IQ > 75, were fluent in English, were able to perform the study tasks, had no previous head injury or prolonged unconsciousness, substance or alcohol dependence, migraine treatments, or MRI contraindications. Additional details about this dataset can be found in previous articles (Keator et al., 2016; Turner et al., 2013). A subset of this dataset was utilized in the current study, summarized in **Table S1**.

**Table S1 |** **FBIRN** **Dataset Summary.** Demographic, clinical characteristics, and cognitive assessments for the FBIRN subset included in the current study are displayed. Participants are assigned to either the schizophrenia or control group. The table displays the number of participants in each group, the range, mean, and standard deviation (SD) of age in years (yrs), and the race distribution. The “Other” category for race includes participants who indicated that they belonged to a race other than those presented, indicated that they belonged to more than one race, or chose not to disclose this information. The following clinical characteristics are reported for the schizophrenia sample: the percentage of participants with medication dosage information available, the mean chlorpromazine (CPZ) equivalent estimate in milligrams (mg) per day, the percentage of participants with age of onset (AO) and duration of illness (DOI) information available, the age of onset and duration of illness, percentage of participants with PANSS Positive (Pos.)/Negative (Neg.)/General (Gen.)/Total scores available, and the mean and SD for each. The following cognitive assessment scores are reported for all participants: the percentage of participants in each group (and total) with Working Memory (WrkMem), Processing Speed (ProcSpd), Verbal Learning (VerbL), Reasoning and Problem Solving (R&PS; Reasoning & Prob. Solv.), Visual Learning (VisL), Attention/Vigilance (AttVig), and Social Cognition (SoCog) domain scores, as well as the mean and SD of the z-scores for each test.

| Dataset | FBIRN | | |
| --- | --- | --- | --- |
|  | **Control** | **Schizophrenia** | **Total** |
|  | Mean ± SD | Mean ± SD | Mean ± SD |
| Demographics |  |  |  |
| N (Male/Female) | 171 (119/52) | 160 (118/42) | 331 (237/94) |
| Age (Range: 18-60 yrs) | 37.13 ± 11.15 | 39.29 ± 11.39 | 38.18 ± 11.30 |
| Race (%) |  |  |  |
| White | 73.68% | 61.25% | 67.67% |
| Black | 12.87% | 23.13% | 17.82% |
| Asian | 9.94% | 13.13% | 11.48% |
| Other | 3.51% | 2.50% | 3.02% |
| Medication |  |  |  |
| Participants w/ CPZ (%) | --- | 77.50% | 77.50% |
| CPZ equivalents (mg) | --- | 354.82 ± 266.47 | --- |
| Symptom Onset & Severity |  |  |  |
| Participants w/ AO & DOI (%) | --- | 98.75% | --- |
| Age of Onset (yrs) | --- | 21.79 ± 7.52 | --- |
| Duration of Illness (yrs) | --- | 17.59 ± 11.12 | --- |
| Participants w/ PANSS Positive (%) | --- | 97.50% | --- |
| PANSS Positive | --- | 15.45 ± 5.22 | --- |
| Participants w/ PANSS Negative (%) | --- | 97.50% | --- |
| PANSS Negative | --- | 14.52 ± 5.73 | --- |
| Participants w/ PANSS General (%) | --- | 97.50% | --- |
| PANSS General | --- | 28.70 ± 7.61 | --- |
| Participants w/ PANSS Total (%) | --- | 97.50% | --- |
| PANSS Total | --- | 58.67 ± 15.54 | --- |
| Cognitive Domain Scores |  |  |  |
| Participants w/ WrkMem (%) | 86.55% | 90.00% | 88.22% |
| Working Memory z-score | 0.03 ± 0.99 | -1.26 ± 1.12 | -0.61 ± 1.24 |
| Participants w/ ProcSpd (%) | 85.96% | 90.00% | 87.92% |
| Processing Speed z-score | -0.00 ± 1.03 | -1.29 ± 1.06 | -0.64 ± 1.23 |
| Participants w/ VerbL (%) | 86.55% | 90.00% | 88.22% |
| Verbal Learning z-score | 0.08 ± 0.96 | -1.33 ± 1.17 | -0.62 ± 1.28 |
| Participants w/ R&PS (%) | 85.96% | 90.00% | 87.92% |
| Reasoning & Problem Solving z-score | 0.03 ± 0.98 | -0.93 ± 1.26 | -0.44 ± 1.22 |
| Participants w/ VisL (%) | 84.80% | 89.38% | 87.01% |
| Visual Learning z-score | 0.00 ± 1.00 | -1.13 ± 1.10 | -0.56 ± 1.19 |
| Participants w/ AttVig (%) | 85.96% | 86.88% | 86.40% |
| Attention/Vigilance z-score | -0.02 ± 1.01 | -1.43 ± 1.44 | -0.70 ± 1.42 |
| Participants w/ SoCog (%) | --- | --- | --- |
| Social Cognition z-score | --- | --- | --- |

COBRE**:** The Center for Biomedical Research Excellence (COBRE; Aine et al., 2017) dataset is comprised of rsfMRI collected from a single site with approval from their IRB. The rsfMRI data was collected using a Siemens 3-Tesla TIM Trio scanner with a standard echo-planar imaging sequence featuring a TR/TE of 2000/29 ms, a voxel size of 3.75 × 3.75 × 4.5 mm with a 1.05 mm slice gap, a field of view of 240 × 240 mm, and a total of 149 volumes. Individuals with schizophrenia were recruited from the University of New Mexico Psychiatric Center, the Raymond G. Murphy Veterans Affairs Medical Center, and from other psychiatric clinics in the Albuquerque metropolitan area and they completed the Structured Clinical Interview for DSM-IV Axis I Disorders for diagnostic confirmation (First et al., 1997). Participants with schizophrenia were excluded if they were not clinically stable. Participants in the control group were recruited through advertisement in the same geographic location. Participants in the control group were excluded if they had a current or past history of a psychiatric disorder (with the exception of one lifetime major depressive episode), had depression or antidepressant use within the past 6 months, lifetime antidepressant use of more than one year, or history of a psychotic disorder in a first-degree relative. All participants gave written informed consent, and were between the ages of 18-65 years old, had no major neurological illnesses, had no previous head injury or prolonged unconsciousness, had no diagnosis of intellectual disability, and had no substance (except for nicotine) or alcohol dependence or abuse. Additional details about this dataset can be found in the original article (Aine et al., 2017). A subset of this dataset was utilized in the current study, summarized in **Table S2**.

**Table S2** | **COBRE** **Dataset Summary.** Demographic, clinical characteristics, and cognitive assessments for the COBRE subset included in the current study are displayed. Participants are assigned to either the schizophrenia or control group. The table displays the number of participants in each group, the range, mean, and standard deviation (SD) of age in years (yrs), and the race distribution. The “Other” category for race includes participants who indicated that they belonged to a race other than those presented, indicated that they belonged to more than one race, or chose not to disclose this information. The following clinical characteristics are reported for the schizophrenia sample: the percentage of participants with medication dosage information available, the mean chlorpromazine (CPZ) equivalent estimate in milligrams (mg) per day, the percentage of participants with age of onset (AO) and duration of illness (DOI) information available, the age of onset and duration of illness, percentage of participants with PANSS Positive (Pos.)/Negative (Neg.)/General (Gen.)/Total scores available, and the mean and SD for each. The following cognitive assessment scores are reported for all participants: the percentage of participants in each group (and total) with Working Memory (WrkMem), Processing Speed (ProcSpd), Verbal Learning (VerbL), Reasoning and Problem Solving (R&PS; Reasoning & Prob. Solv.), Visual Learning (VisL), Attention/Vigilance (AttVig), and Social Cognition (SoCog) domain scores, as well as the mean and SD of the z-scores for each test.

| Dataset | COBRE | | |
| --- | --- | --- | --- |
|  | **Control** | **Schizophrenia** | **Total** |
|  | Mean ± SD | Mean ± SD | Mean ± SD |
| Demographics |  |  |  |
| N (Male/Female) | 89 (64/25) | 78 (64/14) | 167 (128/39) |
| Age (Range: 18-65 yrs) | 38.20 ± 11.54 | 37.36 ± 14.12 | 37.81 ± 12.77 |
| Race (%) |  |  |  |
| White | 89.89% | 85.90% | 88.02% |
| Black | 7.87% | 8.97% | 8.38% |
| Asian | 0 | 0 | 0 |
| Other | 2.25% | 5.13% | 3.59% |
| Medication |  |  |  |
| Participants w/ CPZ (%) | --- | 93.59% | 93.59% |
| CPZ equivalents (mg) | --- | 383.40 ± 306.65 | --- |
| Symptom Onset & Severity |  |  |  |
| Participants w/ AO & DOI (%) | --- | 75.64% | --- |
| Age of Onset (yrs) | --- | 21.05 ± 8.47 | --- |
| Duration of Illness (yrs) | --- | 15.51 ± 12.87 | --- |
| Participants w/ PANSS Positive (%) | --- | 100% | --- |
| PANSS Positive | --- | 15.03 ± 4.91 | --- |
| Participants w/ PANSS Negative (%) | --- | 100% | --- |
| PANSS Negative | --- | 15.04 ± 5.38 | --- |
| Participants w/ PANSS General (%) | --- | 100% | --- |
| PANSS General | --- | 28.58 ± 8.67 | --- |
| Participants w/ PANSS Total (%) | --- | 100% | --- |
| PANSS Total | --- | 58.60 ± 15.01 | --- |
| Cognitive Domain Scores |  |  |  |
| Participants w/ WrkMem (%) | 92.13% | 93.59% | 92.81% |
| Working Memory z-score | 0.43 ± 0.83 | -0.34 ± 0.99 | 0.07 ± 0.98 |
| Participants w/ ProcSpd (%) | 92.13% | 93.59% | 92.81% |
| Processing Speed z-score | 0.71 ± 0.62 | -0.63 ± 0.85 | 0.08 ± 0.99 |
| Participants w/ VerbL (%) | 92.13% | 93.59% | 92.81% |
| Verbal Learning z-score | 0.41 ± 0.91 | -0.36 ± 0.92 | 0.05 ± 0.99 |
| Participants w/ R&PS (%) | 92.13% | 93.59% | 92.81% |
| Reasoning & Problem Solving z-score | 0.55 ± 0.74 | -0.48 ± 0.96 | 0.06 ± 0.99 |
| Participants w/ VisL (%) | 92.13% | 93.59% | 92.81% |
| Visual Learning z-score | 0.37 ± 0.88 | -0.49 ± 1.02 | -0.03 ± 1.04 |
| Participants w/ AttVig (%) | 92.13% | 93.59% | 92.81% |
| Attention/Vigilance z-score | 0.53 ± 0.68 | -0.45 ± 1.04 | 0.07 ± 0.99 |
| Participants w/ SoCog (%) | 92.13% | 93.59% | 92.81% |
| Social Cognition z-score | 0.39 ± 0.84 | -0.49 ± 1.00 | -0.02 ± 1.01 |

MPRC**:** The Maryland Psychiatric Research Center (MPRC; Adhikari et al., 2019) dataset is comprised of rsfMRI collected from three sites, each with approval from their respective IRB. The first site used a Siemens 3-Tesla Allegra scanner and a standard echo-planar imaging sequence with a TR/TE of 2000/27 ms, a voxel size of 3.44 × 3.44 × 4 mm, a field of view of 220 × 220 mm, and 150 volumes. The second site used a Siemens 3-Tesla Trio scanner with a standard echo-planar imaging sequence, a TR/TE of 2210/30 ms, a voxel size of 3.44 × 3.44 × 4 mm, a field of view measuring 220 × 220 mm, and 140 volumes. The third site employed a Siemens 3-Tesla Tim Trio scanner with a standard echo-planar imaging sequence, a TR/TE of 2000/30 ms, a voxel size of 1.72 × 1.72 × 4 mm, a field of view measuring 220 × 220 mm, and 444 volumes. Individuals with schizophrenia were recruited from the outpatient clinics at the MPRC and mental health clinics in the greater Baltimore area and completed the Structured Clinical Interview for DSM-IV Axis I Disorders for diagnostic confirmation (First et al., 1997). Participants in the control group were recruited through media advertisements from the same geographic area. Participants in the control group were excluded if they had a current or past history of a major psychiatric illness or had a family history of psychosis in the prior two generations. All participants gave written informed consent and were between the ages of 10-79 years old, had no major medical or neurological illnesses, history of head injury with cognitive sequelae, diagnosis of intellectual disability, substance abuse in the past 3 months (except for nicotine), alcohol dependence in the past 6 months, or MRI contraindications. Additional details about this dataset can be found in previous articles (Adhikari et al., 2019; Kochunov et al., 2017). A subset of this dataset was utilized in the current study, summarized in **Table S3**.

**Table S3** | **MPRC** **Dataset Summary.** Demographic, clinical characteristics, and cognitive assessments for the MPRC subset included in the current study are displayed. Participants are assigned to either the schizophrenia or control group. The table displays the number of participants in each group, the range, mean, and standard deviation (SD) of age in years (yrs), and the race distribution. The “Other” category for race includes participants who indicated that they belonged to a race other than those presented, indicated that they belonged to more than one race, or chose not to disclose this information. The following clinical characteristics are reported for the schizophrenia sample: the percentage of participants with medication dosage information available, the mean chlorpromazine (CPZ) equivalent estimate in milligrams (mg) per day, the percentage of participants with age of onset (AO) and duration of illness (DOI) information available, the age of onset and duration of illness, percentage of participants with PANSS Positive (Pos.)/Negative (Neg.)/General (Gen.)/Total scores available, and the mean and SD for each. The following cognitive assessment scores are reported for all participants: the percentage of participants in each group (and total) with Working Memory (WrkMem), Processing Speed (ProcSpd), Verbal Learning (VerbL), Reasoning and Problem Solving (R&PS; Reasoning & Prob. Solv.), Visual Learning (VisL), Attention/Vigilance (AttVig), and Social Cognition (SoCog) domain scores, as well as the mean and SD of the z-scores for each test. ^a^BPRS total scores were converted to PANSS total scores for the analysis following Leucht et al. (2013) conversion tables.

| Dataset | MPRC | | |
| --- | --- | --- | --- |
|  | **Control** | **Schizophrenia** | **Total** |
|  | Mean ± SD | Mean ± SD | Mean ± SD |
| Demographics |  |  |  |
| N (Male/Female) | 242 (92/150) | 151 (99/52) | 393 (191/202) |
| Age (Range: 10-79 yrs) | 40.19 ± 15.54 | 38.10 ± 13.64 | 39.38 ± 14.86 |
| Race (%) |  |  |  |
| White | 66.53% | 52.98% | 61.32% |
| Black | 27.69% | 42.38% | 33.33% |
| Asian | 2.89% | 1.99% | 2.54% |
| Other | 2.89% | 2.65% | 2.80% |
| Medication |  |  |  |
| Participants w/ CPZ (%) | 0.83% | 28.48% | 22.52% |
| CPZ equivalents (mg) | --- | 306.74 ± 255.69 | --- |
| Symptom Onset & Severity |  |  |  |
| Participants w/ AO & DOI (%) | --- | --- | --- |
| Age of Onset (yrs) | --- | --- | --- |
| Duration of Illness (yrs) | --- | --- | --- |
| Participants w/ PANSS Positive (%) | --- | --- | --- |
| PANSS Positive | --- | --- | --- |
| Participants w/ PANSS Negative (%) | --- | --- | --- |
| PANSS Negative | --- | --- | --- |
| Participants w/ PANSS General (%) | --- | --- | --- |
| PANSS General | --- | --- | --- |
| Participants w/ PANSS Total (%) | --- | 87.42% | --- |
| PANSS Total^a^ | --- | 66.53 ± 19.24 | --- |
| Cognitive Domain Scores |  |  |  |
| Participants w/ WrkMem (%) | 52.48% | 52.32% | 52.42% |
| Working Memory z-score | 0.47 ± 0.88 | -0.30 ± 0.89 | 0.18 ± 0.96 |
| Participants w/ ProcSpd (%) | 53.72% | 52.98% | 53.44% |
| Processing Speed z-score | 0.51 ± 0.77 | -0.42 ± 0.70 | 0.15 ± 0.87 |
| Participants w/ VerbL (%) | --- | --- | --- |
| Verbal Learning z-score | --- | --- | --- |
| Participants w/ R&PS (%) | --- | --- | --- |
| Reasoning & Problem Solving z-score | --- | --- | --- |
| Participants w/ VisL (%) | --- | --- | --- |
| Visual Learning z-score | --- | --- | --- |
| Participants w/ AttVig (%) | --- | --- | --- |
| Attention/Vigilance z-score | --- | --- | --- |
| Participants w/ SoCog (%) | --- | --- | --- |
| Social Cognition z-score | --- | --- | --- |

B-SNIP 1**:** The Bipolar-Schizophrenia Network on Intermediate Phenotypes 1 (B-SNIP 1; Tamminga et al., 2013) dataset is comprised of rsfMRI collected from six sites (see scanner parameters in **Table S4**), each with approval from their respective IRB. Individuals with schizophrenia were recruited in Baltimore, Chicago, Dallas, Detroit and Boston, and Hartford through referrals from mental health providers and through advertisements and talks at community organizations and support groups. Individuals with schizophrenia completed the Structured Clinical Interview for DSM-IV Axis I Disorders (First et al., 1997) for diagnostic confirmation and were excluded if they were not clinically stable. Participants in the control group were recruited through newspaper and community advertising in the same geographic locations. Participants in the control group were excluded if they had a personal history of a psychiatric disorder or recurrent depression, or had a known immediate family history of psychotic disorders. All participants gave written informed consent and were between the ages of 15-64 years old, had an age-corrected Wide-Range Achievement Test, 4^th^ edition, reading test standard score > 65, and were fluent in English. All participants had no major neurological disorders, history of seizures or head injury with loss of consciousness, substance abuse in the past 30 days, or substance dependence in the past 6 months. Additional details about this dataset can be found in previous articles (Hill et al., 2013; Meda et al., 2014; Tamminga et al., 2013, 2014). A subset of this dataset was utilized in the current study, summarized in **Table S5**.

**Table S4** | **B-SNIP 1** **Scanning Parameters.** Detailed information is provided below for each of the five imaging sites in the B-SNIP 1 dataset.

| **Site** | **TR (ms)** | **Slices (N)** | **Vendor** |
| --- | --- | --- | --- |
| **Baltimore** | 2210 | 36 | Siemens Triotim |
| **Hartford** | 1500 | 29 | Siemens Allegra |
| **Detroit** | 1570 | 29 | Siemens TrioTim |
| **Boston** | 3000 | 30 | GE HDxt |
| **Dallas** | 1500 | 29 | Philips |
| **Chicago** | 1775 | 29 | GE Signa HDX |

**Table S5** | **B-SNIP 1** **Dataset Summary.** Demographic, clinical characteristics, and cognitive assessments for the B-SNIP 1 subset included in the current study are displayed. Participants are assigned to either the schizophrenia or control group. The table displays the number of participants in each group, the range, mean, and standard deviation (SD) of age in years (yrs), and the race distribution. The “Other” category for race includes participants who indicated that they belonged to a race other than those presented, indicated that they belonged to more than one race, or chose not to disclose this information. The following clinical characteristics are reported for the schizophrenia sample: the percentage of participants with medication dosage information available, the mean chlorpromazine (CPZ) equivalent estimate in milligrams (mg) per day, the percentage of participants with age of onset (AO) and duration of illness (DOI) information available, the age of onset and duration of illness, percentage of participants with PANSS Positive (Pos.)/Negative (Neg.)/General (Gen.)/Total scores available, and the mean and SD for each. The following cognitive assessment scores are reported for all participants: the percentage of participants in each group (and total) with Working Memory (WrkMem), Processing Speed (ProcSpd), Verbal Learning (VerbL), Reasoning and Problem Solving (R&PS; Reasoning & Prob. Solv.), Visual Learning (VisL), Attention/Vigilance (AttVig), and Social Cognition (SoCog) domain scores, as well as the mean and SD of the z-scores for each test.

| Dataset | B-SNIP 1 | | |
| --- | --- | --- | --- |
|  | **Control** | **Schizophrenia** | **Total** |
|  | Mean ± SD | Mean ± SD | Mean ± SD |
| Demographics |  |  |  |
| N (Male/Female) | 241 (100/141) | 201 (139/62) | 442 (239/203) |
| Age (Range: 15-64 yrs) | 38.53 ± 12.56 | 34.38 ± 11.95 | 36.64 ± 12.44 |
| Race (%) |  |  |  |
| White | 62.24% | 49.25% | 56.33% |
| Black | 28.63% | 44.28% | 35.75% |
| Asian | 4.98% | 1.99% | 3.62% |
| Other | 4.15% | 4.48% | 4.30% |
| Medication |  |  |  |
| Participants w/ CPZ (%) | --- | 33.33% | 26.37% |
| CPZ equivalents (mg) | --- | 351.92 ± 355.87 | --- |
| Symptom Onset & Severity |  |  |  |
| Participants w/ AO & DOI (%) | --- | 66.67% | --- |
| Age of Onset (yrs) | --- | 21.21 ± 6.77 | --- |
| Duration of Illness (yrs) | --- | 11.40 ± 10.69 | --- |
| Participants w/ PANSS Positive (%) | --- | 100% | --- |
| PANSS Positive | --- | 16.48 ± 5.40 | --- |
| Participants w/ PANSS Negative (%) | --- | 100% | --- |
| PANSS Negative | --- | 16.33 ± 5.89 | --- |
| Participants w/ PANSS General (%) | --- | 100% | --- |
| PANSS General | --- | 31.65 ± 8.65 | --- |
| Participants w/ PANSS Total (%) | --- | 100% | --- |
| PANSS Total | --- | 64.44 ± 16.83 | --- |
| Cognitive Domain Scores |  |  |  |
| Participants w/ WrkMem (%) | 97.51% | 94.53% | 96.15% |
| Working Memory z-score | -0.08 ± 1.11 | -1.26 ± 1.15 | -0.61 ± 1.27 |
| Participants w/ ProcSpd (%) | 97.51% | 95.02% | 96.38% |
| Processing Speed z-score | -0.04 ± 0.98 | -1.36 ± 1.08 | -0.63 ± 1.21 |
| Participants w/ VerbL (%) | 97.51% | 94.53% | 96.15% |
| Verbal Learning z-score | -0.04 ± 1.07 | -1.07 ± 1.33 | -0.50 ± 1.30 |
| Participants w/ R&PS (%) | 97.51% | 94.53% | 96.15% |
| Reasoning & Problem Solving z-score | 0.03 ± 1.00 | -0.53 ± 1.12 | -0.22 ± 1.09 |
| Participants w/ VisL (%) | --- | --- | --- |
| Visual Learning z-score | --- | --- | --- |
| Participants w/ AttVig (%) | --- | --- | --- |
| Attention/Vigilance z-score | --- | --- | --- |
| Participants w/ SoCog (%) | --- | --- | --- |
| Social Cognition z-score | --- | --- | --- |

B-SNIP 2**:** The Bipolar-Schizophrenia Network on Intermediate Phenotypes 2 (B-SNIP 2; Clementz et al., 2022) dataset is comprised of rsfMRI collected from five sites (see scanner parameters in **Table S6**), each with approval from their respective IRB. Individuals were recruited in Georgia, Hartford, Boston, Dallas, and Chicago following the same recruitment procedures utilized for B-SNIP 1 (see Tamminga et al., 2013). Individuals with schizophrenia completed the Structured Clinical Interview for DSM-IV Axis I Disorders (First et al., 1997), were diagnosed according to the DSM-IV-TR, and were on stable medication with no major changes in the past 30 days. Participants in the control group were excluded if they had a personal history of a psychotic disorder or recurrent mood disorder, or had a known close relative with these disorders. All participants gave written informed consent and were between the ages of 18-65 years old, had an age-corrected Wide-Range Achievement Test, 4th edition, reading test standard score > 65, and were fluent in English. All participants had no history of head injury with loss of consciousness > 10 min, no diagnosis of substance abuse in the past 30 days, no substance dependence in the past 3 months, no positive urine toxicology for drugs of abuse on the day of testing, no history of systemic medical or neurological disorder affecting mood or cognition, and no intellectual disability. Additional details about this dataset can be found in previous articles (Clementz et al., 2022; Meda et al., 2025; Ward et al., 2025). A subset of this dataset was utilized in the current study, summarized in **Table S7**.

**Table S6** | **B-SNIP 2** **Scanning Parameters.** Detailed information is provided below for each of the five imaging sites in the B-SNIP 2 dataset.

| **Site** | **TR (ms)** | **TE (ms)** | **Slices (N)** | **Acquisition Matrix (mm)** | **Voxel Size (mm)** | **Vendor** |
| --- | --- | --- | --- | --- | --- | --- |
| **Georgia** | 2000 | 30 | 30 | 64x64 | 3.4x3.4x5 | GE HDx |
| **Hartford** | 2000 | 30 | 30 | 64x64 | 3.4x3.4x5 | Siemens Skyra |
| **Boston** | 2000 | 30 | 30 | 64x64 | 3.4x3.4x4 | GE HDxt |
| **Dallas** | 2000 | 30 | 30 | 64x64 | 3.4x3.4x4 | Philips Achieva |
| **Chicago** | 2000 | 30 | 30 | 64x64 | 3.4x3.4x4 | Philips dStream Achieva |

**Table S7** | **B-SNIP 2** **Dataset Summary.** Demographic, clinical characteristics, and cognitive assessments for the B-SNIP 2 subset included in the current study are displayed. Participants are assigned to either the schizophrenia or control group. The table displays the number of participants in each group, the range, mean, and standard deviation (SD) of age in years (yrs), and the race distribution. The “Other” category for race includes participants who indicated that they belonged to a race other than those presented, indicated that they belonged to more than one race, or chose not to disclose this information. The following clinical characteristics are reported for the schizophrenia sample: the percentage of participants with medication dosage information available, the mean chlorpromazine (CPZ) equivalent estimate in milligrams (mg) per day, the percentage of participants with age of onset (AO) and duration of illness (DOI) information available, the age of onset and duration of illness, percentage of participants with PANSS Positive (Pos.)/Negative (Neg.)/General (Gen.)/Total scores available, and the mean and SD for each. The following cognitive assessment scores are reported for all participants: the percentage of participants in each group (and total) with Working Memory (WrkMem), Processing Speed (ProcSpd), Verbal Learning (VerbL), Reasoning and Problem Solving (R&PS; Reasoning & Prob. Solv.), Visual Learning (VisL), Attention/Vigilance (AttVig), and Social Cognition (SoCog) domain scores, as well as the mean and SD of the z-scores for each test.

| Dataset | B-SNIP 2 | | |
| --- | --- | --- | --- |
|  | **Control** | **Schizophrenia** | **Total** |
|  | Mean ± SD | Mean ± SD | Mean ± SD |
| Demographics |  |  |  |
| N (Male/Female) | 398 (161/237) | 299 (182/117) | 697 (343/354) |
| Age (Range: 18-60 yrs) | 34.13 ± 11.81 | 38.00 ± 11.97 | 35.79 ± 12.02 |
| Race (%) |  |  |  |
| White | 49.75% | 36.12% | 43.90% |
| Black | 33.92% | 49.83% | 40.75% |
| Asian | 7.79% | 4.35% | 6.31% |
| Other | 8.54% | 9.70% | 9.04% |
| Medication |  |  |  |
| Participants w/ CPZ (%) | --- | 60.87% | 60.87% |
| CPZ equivalents (mg) | --- | 432.32 ± 368.30 | --- |
| Symptom Onset & Severity |  |  |  |
| Participants w/ AO & DOI (%) | --- | 95.32% | --- |
| Age of Onset (yrs) | --- | 20.04 ± 8.11 | --- |
| Duration of Illness (yrs) | --- | 18.21 ± 12.34 | --- |
| Participants w/ PANSS Positive (%) | --- | 99.67% | --- |
| PANSS Positive | --- | 16.30 ± 6.17 | --- |
| Participants w/ PANSS Negative (%) | --- | 99.67% | --- |
| PANSS Negative | --- | 16.01 ± 6.55 | --- |
| Participants w/ PANSS General (%) | --- | 99.33% | --- |
| PANSS General | --- | 30.09 ± 9.34 | --- |
| Participants w/ PANSS Total (%) | --- | 99.33% | --- |
| PANSS Total | --- | 62.48 ± 19.18 | --- |
| Cognitive Domain Scores |  |  |  |
| Participants w/ WrkMem (%) | 97.99% | 94.31% | 96.41% |
| Working Memory z-score | -0.31 ± 1.01 | -1.30 ± 1.29 | -0.73 ± 1.23 |
| Participants w/ ProcSpd (%) | 97.99% | 93.98% | 96.27% |
| Processing Speed z-score | -0.18 ± 1.06 | -1.34 ± 1.15 | -0.67 ± 1.24 |
| Participants w/ VerbL (%) | 97.74% | 94.31% | 96.27% |
| Verbal Learning z-score | -0.28 ± 1.25 | -1.41 ± 1.39 | -0.76 ± 1.42 |
| Participants w/ R&PS (%) | 97.99% | 94.31% | 96.41% |
| Reasoning & Problem Solving z-score | 0.03 ± 0.92 | -0.61 ± 1.09 | -0.24 ± 1.04 |
| Participants w/ VisL (%) | --- | --- | --- |
| Visual Learning z-score | --- | --- | --- |
| Participants w/ AttVig (%) | --- | --- | --- |
| Attention/Vigilance z-score | --- | --- | --- |
| Participants w/ SoCog (%) | --- | --- | --- |
| Social Cognition z-score | --- | --- | --- |

Dataset 6**:** The final dataset is comprised of rsfMRI collected from four imaging sites in China (see scanner parameters in **Table S8**), each with approval from their respective research ethics board. Individuals with schizophrenia were recruited from Anhui Medical University (AH), Beijing Huilongguan Hospital (BJHLG), Guangzhou Psychiatric Hospital (GZ), and Zhumadian Psychiatric Hospital (ZMD) and they completed the Structured Clinical Interview for DSM-IV-TR Axis I Disorders for diagnostic confirmation. Participants in the control group were recruited through advertisements in the same geographic location and were excluded if they had a psychiatric disorder. All participants gave written informed consent, and were between the ages of 16-54 years old and were right-handed and Chinese Han ethnicity. All participants had no major neurological illnesses, previous head injury or prolonged unconsciousness, prior electroconvulsive therapy, were not pregnant, and had no substance or alcohol dependence or abuse. Additional details about this dataset can be found in previous articles (Meng et al., 2023; Yan et al., 2019, 2024). A subset of this dataset was utilized in the current study, summarized in **Table S9**.

**Table S8** | **Dataset 6** **Scanning Parameters.** Detailed information is provided below for the four imaging sites utilized from dataset 6.

| **Site** | **TR (ms)** | **TE (ms)** | **Slices (N)** | **Acquisition Matrix (mm)** | **Voxel Size (mm)** | **Vendor** |
| --- | --- | --- | --- | --- | --- | --- |
| **AH** | 2000 | 30 | 33 | 64x64 | 3.44×3.44×4.60 | GE Signa HDxt 3T |
| **BJHLG** | 2000 | 30 | 33 | 64x64 | 3.44×3.44×4.60 | Siemens TrioTim 3T |
| **GZ** | 2000 | 30 | 33 | 64x64 | 3.44×3.44×4.60 | Phillips Achieva 3T |
| **ZMD** | 2000 | 30 | 33 | 64x64 | 3.44×3.44×4.60 | GE Signa HDxt 3T |

**Table S9** | **Dataset 6 Summary.** Demographic, clinical characteristics, and cognitive assessments for the dataset 6 subset included in the current study are displayed. Participants are assigned to either the schizophrenia or control group. The table displays the number of participants in each group, the range, mean, and standard deviation (SD) of age in years (yrs), and the race distribution. The “Other” category for race includes participants who indicated that they belonged to a race other than those presented, indicated that they belonged to more than one race, or chose not to disclose this information. The following clinical characteristics are reported for the schizophrenia sample: the percentage of participants with medication dosage information available, the mean chlorpromazine (CPZ) equivalent estimate in milligrams (mg) per day, the percentage of participants with age of onset (AO) and duration of illness (DOI) information available, the age of onset and duration of illness, percentage of participants with PANSS Positive (Pos.)/Negative (Neg.)/General (Gen.)/Total scores available, and the mean and SD for each. The following cognitive assessment scores are reported for all participants: the percentage of participants in each group (and total) with Working Memory (WrkMem), Processing Speed (ProcSpd), Verbal Learning (VerbL), Reasoning and Problem Solving (R&PS; Reasoning & Prob. Solv.), Visual Learning (VisL), Attention/Vigilance (AttVig), and Social Cognition (SoCog) domain scores, as well as the mean and SD of the z-scores for each test.

| Dataset | Dataset 6 | | |
| --- | --- | --- | --- |
|  | **Control** | **Schizophrenia** | **Total** |
|  | Mean ± SD | Mean ± SD | Mean ± SD |
| Demographics |  |  |  |
| N (Male/Female) | 267 (143/124) | 359 (189/170) | 626 (332/294) |
| Age (Range: 16-54 yrs) | 28.06 ± 7.44 | 29.08 ± 7.82 | 28.65 ± 7.67 |
| Race (%) |  |  |  |
| White | 0 | 0 | 0 |
| Black | 0 | 0 | 0 |
| Asian | 100.00% | 100.00% | 100.00% |
| Other | 0 | 0 | 0 |
| Medication |  |  |  |
| Participants w/ CPZ (%) | --- | 20.33% | 20.33% |
| CPZ equivalents (mg) | --- | 399.32 ± 166.20 | --- |
| Symptom Onset & Severity |  |  |  |
| Participants w/ AO & DOI (%) | --- | 43.45% | --- |
| Age of Onset (yrs) | --- | 24.41 ± 7.13 | --- |
| Duration of Illness (yrs) | --- | 5.23 ± 5.10 | --- |
| Participants w/ PANSS Positive (%) | --- | 99.72% | --- |
| PANSS Positive | --- | 24.27 ± 4.01 | --- |
| Participants w/ PANSS Negative (%) | --- | 99.72% | --- |
| PANSS Negative | --- | 18.54 ± 6.46 | --- |
| Participants w/ PANSS General (%) | --- | 99.72% | --- |
| PANSS General | --- | 38.06 ± 7.42 | --- |
| Participants w/ PANSS Total (%) | --- | 99.72% | --- |
| PANSS Total | --- | 80.47 ± 14.08 | --- |
| Cognitive Domain Scores |  |  |  |
| Participants w/ WrkMem (%) | 98.50% | 87.74% | 92.33% |
| Working Memory z-score | 0.33 ± 0.87 | -0.27 ± 0.98 | 0.00 ± 0.98 |
| Participants w/ ProcSpd (%) | --- | --- | --- |
| Processing Speed z-score | --- | --- | --- |
| Participants w/ VerbL (%) | --- | --- | --- |
| Verbal Learning z-score | --- | --- | --- |
| Participants w/ R&PS (%) | --- | --- | --- |
| Reasoning & Problem Solving z-score | --- | --- | --- |
| Participants w/ VisL (%) | --- | --- | --- |
| Visual Learning z-score | --- | --- | --- |
| Participants w/ AttVig (%) | --- | --- | --- |
| Attention/Vigilance z-score | --- | --- | --- |
| Participants w/ SoCog (%) | --- | --- | --- |
| Social Cognition z-score | --- | --- | --- |

Supplementary Information S2. Cognitive Assessment Harmonization

The current study utilized the seven cognitive domains of the Measurement and Treatment Research to Improve Cognition in Schizophrenia (MATRICS; Green & Nuechterlein, 2004) Consensus Cognitive Battery (MCCB; Nuechterlein et al., 2008) to assess cognitive performance. These include working memory, speed of processing, verbal learning, reasoning and problem solving, visual learning, attention/vigilance, and social cognition. Although assessments varied across datasets, the psychometric tests were comparable within each domain (see **Table S10** for a comparison chart) and the scores were standardized across datasets. To standardize each domain measure, cognitive scores were z-scored independently for each dataset so that each cognitive domain score represented cognitive performance relative to other participants within the same study (via the same measurement approach).

To further elaborate on the comparability of the tests within each cognitive domain, we note the following: The COBRE dataset utilized all seven domains of the MCCB (Aine et al., 2017). The FBIRN dataset utilized the Computerized Multiphasic Interactive Neurocognitive System (CMINDS), which is a computerized version of the MCCB and its comparability is described at length in (Van Erp et al., 2015). For the working memory domain, the B-SNIP and MPRC datasets utilized the Brief Assessment of Cognition in Schizophrenia (BACS) digit sequencing task (DST; Keefe, 2004; Keefe et al., 2008), which is a subset of the working memory domain tests administered in the MCCB and CMINDS. In this task, individuals are presented with clusters of numbers of increasing length and then asked to repeat back to the test administrator the numbers in order (with forward and backward subtasks), and the score reflects the number of trials with all items in the correct order as well as the longest sequence recalled correctly. Dataset 6 utilized a comparable digit span test described in Liu et al. (2019), in which individuals are likewise presented with a series of digits which they must repeat back to the test administrator in the correct order (again with forward and backward subtasks), with the score reflecting the maximum number of digits repeated correctly. For the speed of processing domain, the category fluency and symbol coding subtests were identical across COBRE, FBIRN, and B-SNIP datasets, although COBRE and FBIRN included the trail making test (Nuechterlein et al., 2008), while B-SNIP used the token motor task (Keefe, 2004; Keefe et al., 2008) instead (both of which measure motor speed). The MPRC dataset utilized the symbol coding subtest (Wechsler, 1997), a subset of the measures included for the speed of processing domain used in the other datasets. For the verbal learning domain, the MCCB, CMINDS and BACS list learning tests were used for COBRE, FBIRN, and B-SNIP datasets respectively. In all tests, episodic memory functions are measured as individuals are read a word list and then asked to recall as many words as possible. For the reasoning and problem solving domain, the FBIRN dataset utilized a maze test (a subtest with the MCCB reasoning and problem solving domain), where individuals are presented with mazes increasing in difficulty and scored on accuracy and speed. Alternatively, the B-SNIP datasets utilized the BACS tower of London subtest (also a subset within the MCCB reasoning and problem solving domain), in which individuals are presented with two pictures simultaneously with differing arrangements of colored balls and asked to give the number of times the balls in one picture need to be moved to match the other picture. The score is based on accuracy over 20 trials with increasing difficulty. Although these tests differ (note that the MCCB used in COBRE is a composite of both), both measure constructs of executive function, planning, and problem-solving.

| **Cognitive Domain** | **COBRE** | **FBIRN** | **B-SNIP 1 & 2** | **MPRC** | **Dataset 6** |
| --- | --- | --- | --- | --- | --- |
| Working  Memory | MCCB  (spatial & LNS) | CMINDS  (spatial & LNS) | BACS: DST  (forward & backward) | BACS: DST  (forward & backward) | Digit Span  (forward & backward) |
| Speed of Processing | MCCB  (trails; cat. fluency; symbol coding) | CMINDS  (trails; cat. fluency; symbol coding) | BACS  (token motor; cat. fluency; symbol coding) | WAIS-III  (symbol coding) | NA |
| Verbal  Learning | MCCB  (list learning) | CMINDS  (list learning) | BACS  (list learning) | NA | NA |
| Reasoning & Problem Solving | MCCB  (mazes; tower of London) | CMINDS  (mazes) | BACS  (tower of London) | NA | NA |
| Visual  Learning | MCCB  (figure learning) | CMINDS  (figure learning) | NA | NA | NA |
| Attention/  Vigilance | MCCB  (cont. performance) | CMINDS  (cont. performance) | NA | NA | NA |
| Social Cognition | MCCB  (manage emotions) | NA | NA | NA | NA |

**Table S10** | **Cognitive Assessments**. The specific cognitive assessments utilized from each dataset for each of the cognitive domains reported in the main text are shown in the table below. MCCB: MATRICS Consensus Cognitive Battery (Nuechterlein et al., 2008); LNS: letter-number sequencing subtest; CMINDS: Computerized Multiphasic Interactive Neurocognitive System (Van Erp et al., 2015); BACS: Brief Assessment of Cognition in Schizophrenia (Keefe, 2004; Keefe et al., 2008); the Dataset 6 Digit Span test is described in detail in Liu et al. (2019) and is comparable to the BACS digit sequencing task (DST); WAIS-III: Wechsler Adult Intelligence Scale, 3rd ed. (Wechsler, 1997), the digit symbol-coding subtest is one of the included measures of the MCCB Speed of Processing domain.

Supplementary Information S3. NeuroMark QC Criteria

The quality control (QC) criteria used in the current study is consistent with NeuroMark QC protocol described in (Du et al., 2020). This includes the following: Head motion movement is below 3º rotations and 3 mm translations in each direction. The mean framewise displacement defined as the average of the sum of absolute instantaneous head motions across the six motion realignment parameters should be less than 0.25. We ensured every subject had a good normalization to the template by comparing individual and group masks. For each subject, we calculated the individual mask by setting voxels in the first fMRI time volume which were greater than 90% of the whole brain mean to 1. We computed group mask by setting voxels which were included in more than 90% of subjects to 1. For each subject, we calculated the spatial correlations (i.e., Pearson correlation) between the group mask and the individual mask. The spatial correlations were calculated using voxels within the top 10 slices of the mask, within the bottom 10 slices of the mask and within the whole mask, resulting in three correlation values for each subject. If a subject had correlations larger than 0.75 for the top 10 slices, larger than 0.55 for the bottom 10 slices, and larger than 0.8 for the whole mask, we included this subject for further analysis. These thresholds have been utilized in previous studies (Du et al., 2020; Jensen et al., 2024; Yan et al., 2024) to ensure a high-quality mask and fMRI data for all individuals.

Supplementary Information S4. QC Validation Results

In each QC validation analysis, we investigated only those FNCs showing significant associations in the primary group comparison analysis. As we corrected for multiple comparisons in the primary analysis and masked subsequent tests, only testing those with FDR corrected significance at the group-level, we did not perform FDR correction on subsequent analyses to avoid over-correcting. We then compared the results between each analysis with the full sample and the more stringent QC subset by performing a Pearson correlation between the results from each as well as by calculating the percentage of significant FNCs in each. Our motivation for performing this QC validation approach as opposed to running analyses with only the reduced QC sample to begin with was that QC thresholds vary greatly across studies and can be somewhat arbitrary. Such thresholds may needlessly reduce the sample size – as well as sample heterogeneity - and in turn result in the analyses achieving reduced statistical power which may be critical for identifying smaller effects (Marek et al., 2022). This is especially a concern in the current analyses, as clinical groups are prone to higher head motion. Our approach acknowledges that overly stringent methods for addressing head motion artifacts may reduce artifactual influence at the cost of losing potentially meaningful signal (Bright & Murphy, 2015; Kumar et al., 2024) and seeks to overcome threshold-driven reporting bias (Amrhein et al., 2019; McShane et al., 2019; Wedel & Gal, 2024) and maximize opportunities to explore potentially meaningful trends in the data while also validating that the results are not driven by head motion artifacts. The results of each QC analysis are reported below.

***QC FNC Group Comparisons***

We repeated the group comparison analysis in a QC subset consisting of 65% of the original sample, with 931 subjects excluded (schizophrenia = 518; control = 413) which did not meet the QC criteria. We found a very strong correlation (*r* = .97) between the effect sizes from the main analysis with the full sample and the QC sample. Furthermore, 86.68% of the significant (*q* < .05) FNC pairs in the main analysis were also significant (*p* < .05) in the QC analysis (see **Figure S1**).

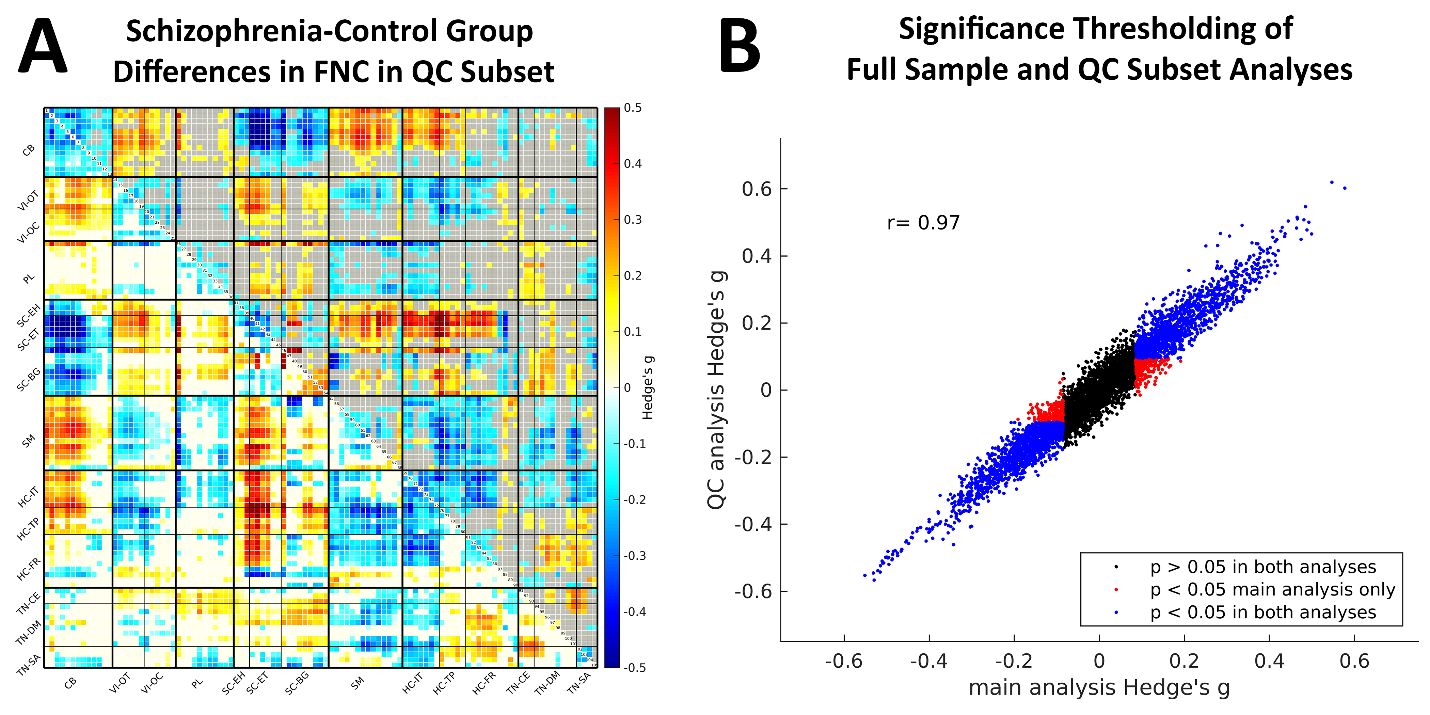

**Figure S1** | **QC Validation and the Effects of Thresholding.** Group comparisons in the quality control (QC) subset for each combination of the 105 NeuroMark intrinsic connectivity networks (ICNs) are displayed in the matrix **(A)**. The upper triangle displays only significant (*p* < .05) case-control group differences in functional network connectivity (FNC) with non-significant pairs grayed out. The heatmap uses red to represent an increase in FNC in schizophrenia relative to controls and blue is used to represent a relative decrease in schizophrenia. ICNs are grouped by their domain-subdomain labels, consistent with the NeuroMark_fMRI_2.2 template: cerebellar (CB), visual-occipitotemporal (VI-OT), visual-occipital (VI-OC), paralimbic (PL), subcortical-extended hippocampal (SC-EH), subcortical-extended thalamic (SC-ET), subcortical-basal ganglia (SC-BG), sensorimotor (SM), higher cognition-insular-temporal (HC-IT), higher cognition-temporoparietal (HC-TP), higher cognition-frontal (HC-FR), triple network-central executive (TN-CE), triple network-default mode (TN-DM), and triple network-salience (TN-SA). **(B)** The effect sizes (Hedge’s g) from the group comparisons with the full sample (N = 2,656) and QC subset (N = 1,725) are displayed in a scatterplot. Case/control group associations with FNC which were significant (*p* < .05) in both analyses are represented in blue, while associations with FNC which were only significant (*p* < .05) in the full sample are represented in red. Group associations with FNC which were not significant (*p* > 0.05) in either analysis are represented in black.

***Antipsychotic Medication and QC FNC***

We repeated the medication association analysis in a reduced QC subset (*N* = 259). We found a strong correlation (*r* = .728) between the results from the main analysis with the full sample and the QC sample. Although, only 34.93% of the FNC pairs which displayed significant (*p* < .05) correlations with CPZ in the full sample were also significantly (*p* < .05) correlated with CPZ in the QC subset.

***Onset, Chronicity, and QC FNC***

We repeated the AO association analysis in a reduced QC subset (N = 416) and found a strong correlation (*r* = .657) between the results from the main analysis with the full sample and the QC sample. In this analysis, 36.19% of the FNC pairs which displayed significant (*p* < .05) correlations with AO in the full sample were also significantly (*p* < .05) correlated with AO in the QC subset. We repeated the DOI association analysis in a reduced QC subset (*N* = 416). We found a strong correlation (*r* = .747) between the results from the main analysis with the full sample and the QC sample. 36.36% of the FNC pairs which displayed significant (*p* < .05) correlations with DOI in the full sample were also significantly (*p* < .05) correlated with DOI in the QC subset.

***Symptom Severity and QC FNC***

We repeated the PANSS association analyses in reduced QC subsets which included 57.7-58.5% of the initial samples for each (see **Table S11**). We observed relatively strong correlations (*r* = .645 - .77) between the results from the main analyses with the full samples and the QC samples. However, most likely due to the reduced sample sizes, relatively few (ranging from 22.41% to 47.20%) of the FNC pairs displaying significant (*p* < .05) correlations with PANSS in the full samples were also significantly (*p* < .05) correlated with PANSS in the QC subsets.

**Table S11** | **FNC-PANSS Associations in the QC Subsets**. The number of participants from the group comparison analysis which had information available for each of the PANSS subscales is reported along with the number retained in the more stringent quality control (QC) subset. Next, the percentage of significant FNC-PANSS associations retained in the QC subset are displayed along with the Pearson correlation coefficient r and accompanying p-value for the correlation between the results of the full and reduced sample FNC-PANSS analyses.

| **Subscale** | **N** | **QC Subset N** | **% significant in QC** | ***r*** | ***p*** |
| --- | --- | --- | --- | --- | --- |
| **PANSS Positive** | 1091 | 630 | 47.20% | 0.748 | < .05 |
| **PANSS Negative** | 1091 | 630 | 22.41% | 0.645 | < .05 |
| **PANSS General** | 1090 | 629 | 29.86% | 0.761 | < .05 |
| **PANSS Total** | 1222 | 715 | 31.56% | 0.77 | < .05 |

***Cognitive Performance and QC FNC***

We repeated each of the cognitive assessment analyses with reduced QC subsets ranging from 36.13% - 65.85% of the initial samples (see **Table S12**). Correlations between the results of the initial and QC samples ranged from moderate to high (r = .47 - .809) with relatively few (< 54%) of the significant (*p* < .05) FNC pairs retained in the reduced QC samples. It appears that the QC validation was largely impacted by the reduced sample sizes, as most of the effects were relatively small to begin with and the samples were reduced quite dramatically. To further demonstrate the effects of reduced sample size, we performed a post-hoc analysis by randomly sampling reduced subsets of equal size with the QC analysis and found similar results whether QC criteria were considered or not (see **Supplementary Information S8**).

**Table S12 | FNC-Cognitive Assessment Associations in the QC Subsets**. The number of participants from the group comparison analysis which had cognitive assessment scores is reported for seven cognitive domains. The number retained in the more stringent quality control (QC) subset is then reported. Next, the percentage of significant FNC-Cognitive score associations retained in the QC subset are displayed along with the Pearson correlation coefficient r and accompanying p-value for the correlation between the results of the full and reduced sample FNC-Cognitive score analyses.

| **MCCB Domain** | **N** | **QC Subset N** | **% significant in QC** | ***r*** | ***p*** |
| --- | --- | --- | --- | --- | --- |
| **Speed of Processing** | 1753 | 1021 | 47.47% | 0.809 | < .05 |
| **Attention/Vigilance** | 441 | 176 | 15.04% | 0.47 | < .05 |
| **Working Memory** | 2328 | 1533 | 33.09% | 0.69 | < .05 |
| **Verbal Learning** | 1543 | 868 | 38.24% | 0.745 | < .05 |
| **Visual Learning** | 443 | 175 | 53.42% | 0.759 | < .05 |
| **Reasoning & Problem Solving** | 1543 | 868 | 31.58% | 0.746 | < .05 |
| **Social Cognition** | 155 | 56 | 28.45% | 0.503 | < .05 |

Supplementary Information S5. Site-Level Analyses

To explore the potential effects of imaging site, the main case-control group difference analysis was repeated with each individual imaging site with the full and quality control (QC) subsets (see **Table S13**). Pearson correlations between the effect sizes (Hedge’s g) of each individual site and the effect sizes of the main analysis with the full dataset demonstrate that the main effects were observed in nearly every imaging site, with the similarity of results scaling with sample size in most cases. In addition, we further investigated consensus across sites utilizing effect size (see **Figure S2**). For each of the functional network connectivity (FNC) features with effect sizes of Hedge’s g >= .2 in the main analysis, the number of sites which also displayed effect sizes of Hedge’s g >= .2 for the given FNC feature was tallied. The majority (78.64%) of FNC features with effect sizes of g >= .2 in the main analysis were consistently identified in at least half (14) of the imaging sites included in the analysis. This finding demonstrates the consistency of these patterns across datasets and sites and suggests that potential site effects were negligible.

**Table S13** | **Site Level Analyses.** To explore the potential effects of imaging site, the main analysis was repeated with each individual site with the full and quality control (QC) subsets. Here we report the results of a Pearson correlation between the results of each individual site and the results of the main analysis with the full dataset.

| **Dataset** - Site | ***N*** | ***r*** | ***p*** | **QC *N*** | **QC *r*** | **QC *p*** |
| --- | --- | --- | --- | --- | --- | --- |
| **B-SNIP 1** |  | | | | | |
| detroit | 45 | 0.253 | < 0.001 | 27 | 0.180 | < 0.001 |
| baltimore | 120 | 0.635 | < 0.001 | 100 | 0.604 | < 0.001 |
| boston_b1 | 13 | 0.118 | < 0.001 | 12 | 0.132 | < 0.001 |
| dallas_b1 | 82 | 0.466 | < 0.001 | 57 | 0.270 | < 0.001 |
| chicago_b1 | 83 | 0.469 | < 0.001 | 65 | 0.468 | < 0.001 |
| hartford_b1 | 99 | 0.427 | < 0.001 | 62 | 0.407 | < 0.001 |
| **B-SNIP 2** |  | | | | | |
| chicago_b2 | 156 | 0.616 | < 0.001 | 92 | 0.531 | < 0.001 |
| dallas_b2 | 115 | 0.607 | < 0.001 | 52 | 0.406 | < 0.001 |
| boston_b2 | 79 | 0.547 | < 0.001 | 46 | 0.508 | < 0.001 |
| hartford_b2 | 198 | 0.689 | < 0.001 | 116 | 0.508 | < 0.001 |
| georgia | 149 | -0.047 | < 0.001 | 87 | 0.096 | < 0.001 |
| **MPRC** |  | | | | | |
| pk_1 | 110 | 0.581 | < 0.001 | 78 | 0.548 | < 0.001 |
| pk_2 | 131 | 0.574 | < 0.001 | 103 | 0.569 | < 0.001 |
| pk_3 | 152 | 0.513 | < 0.001 | 87 | 0.469 | < 0.001 |
| **FBIRN** |  | | | | | |
| fbirn_12 | 30 | 0.338 | < 0.001 | 12 | 0.192 | < 0.001 |
| fbirn_10 | 28 | 0.255 | < 0.001 | 10 | 0.168 | < 0.001 |
| fbirn_13 | 60 | 0.654 | < 0.001 | 25 | 0.311 | < 0.001 |
| fbirn_18 | 60 | 0.671 | < 0.001 | 28 | 0.472 | < 0.001 |
| fbirn_3 | 55 | 0.648 | < 0.001 | 31 | 0.657 | < 0.001 |
| fbirn_7 | 27 | 0.222 | < 0.001 | 7 | NA | NA |
| fbirn_8 | 5 | NA | NA | 4 | NA | NA |
| fbirn_9 | 66 | 0.687 | < 0.001 | 18 | 0.533 | < 0.001 |
| **COBRE** |  | | | | | |
| cobre | 167 | 0.756 | < 0.001 | 59 | 0.540 | < 0.001 |
| **Dataset 6** |  | | | | | |
| AH | 118 | 0.641 | < 0.001 | 91 | 0.640 | < 0.001 |
| BJHLG | 147 | 0.765 | < 0.001 | 121 | 0.671 | < 0.001 |
| GZ | 212 | 0.726 | < 0.001 | 206 | 0.727 | < 0.001 |
| ZMD | 149 | 0.556 | < 0.001 | 129 | 0.516 | < 0.001 |
| **All Sites** | 2656 | 1 | < 0.001 | 1725 | 0.970 | < 0.001 |

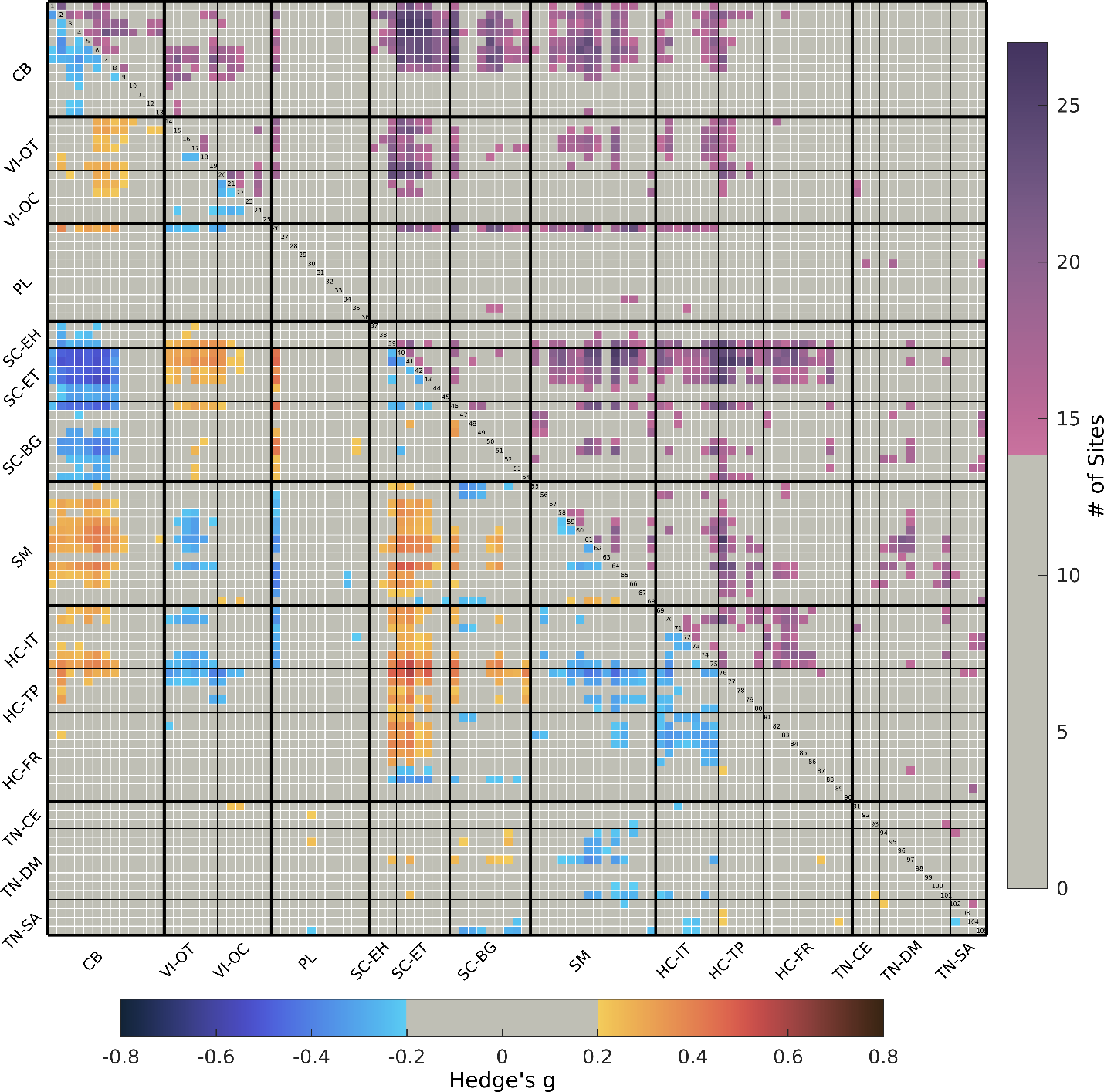

**Figure S2** | **Group Comparison Analysis Site Consensus.** The effect sizes (Hedge’s *g*) of the group differences in the main analysis are displayed in the lower triangle of the matrix. The lower triangle is thresholded to display only FNC features with effect sizes of Hedge’s *g* >= .2. To explore the consistency of group effects in the main analysis across individual imaging sites, for each of the functional network connectivity (FNC) features with effect sizes of Hedge’s *g* >= .2 in the main analysis, the number of sites which also displayed effect sizes of Hedge’s *g* >= .2 for the given FNC feature has been tallied and is displayed in the upper triangle of the matrix. The upper triangle is thresholded to display only FNC features with effect sizes of Hedge’s *g* >= .2 in at least 14 sites. ICNs are grouped by their domain-subdomain labels, consistent with the NeuroMark_fMRI_2.2 template: cerebellar (CB), visual-occipitotemporal (VI-OT), visual-occipital (VI-OC), paralimbic (PL), subcortical-extended hippocampal (SC-EH), subcortical-extended thalamic (SC-ET), subcortical-basal ganglia (SC-BG), sensorimotor (SM), higher cognition-insular-temporal (HC-IT), higher cognition-temporoparietal (HC-TP), higher cognition-frontal (HC-FR), triple network-central executive (TN-CE), triple network-default mode (TN-DM), and triple network-salience (TN-SA).

Supplementary Information S6. Percentage of Group Effects Associated with Medication, Chronicity, Symptom Severity, and Cognitive Performance

**Table S14** | **Percentage of Group Effects Associated with Analysis Variables.** 3,116 of the 5,460 total FNC pairs displayed statistically significant differences in schizophrenia. Each of these was tested in subsequent analyses for associations with medication, chronicity, symptom severity, and cognitive performance. For each analysis, the total number of these FNC pairs showing statistically significant associations with the given variable is reported in the table below accompanied by its corresponding percentage from all FNC pairs with significant group associations.

| **Variable** | **# Significant FNC Pairs** | **% of Sig. FNCs from Group Analysis** |
| --- | --- | --- |
| CPZ | 229 | 7.35% |
| PANSS_pos | 161 | 5.17% |
| PANSS_neg | 241 | 7.73% |
| PANSS_gen | 221 | 7.09% |
| PANSS_total | 225 | 7.22% |
| WM_z | 136 | 4.36% |
| ProcSpd_z | 257 | 8.25% |
| VerbL_z | 272 | 8.73% |
| RProbS_z | 209 | 6.71% |
| VisL_z | 365 | 11.71% |
| AttVig_z | 226 | 7.25% |
| SoCog_z | 116 | 3.72% |
| dur_ill_yrs | 484 | 15.53% |
| AO | 210 | 6.74% |
| **Group** | **3116** | **1** |

Supplementary Information S7. Group Differences Accounting for CPZ

To further investigate the effects of CPZ on the diagnostic effects reported in the main analysis, we performed a general linear model (GLM) with FNC as the response variable, group (schizophrenia/control) as the predictor, and age, sex, race, imaging site, mean framewise displacement, and CPZ as covariates. For the schizophrenia group, a reduced subset of individuals with CPZ scores was utilized (*N* = 562). The control group was assigned CPZ scores of zero. The results of this group comparison analysis controlling for CPZ are presented in **Figure S3**. A Pearson correlation between the effect sizes (Hedge’s g) of the primary group comparison analysis and this analysis controlling for CPZ revealed that the results were highly similar (*r* = .9182, *p* < 0.05), with a reduced number of FNC pairs achieving significance in the current analysis (61.5%), likely due to the reduced sample size.

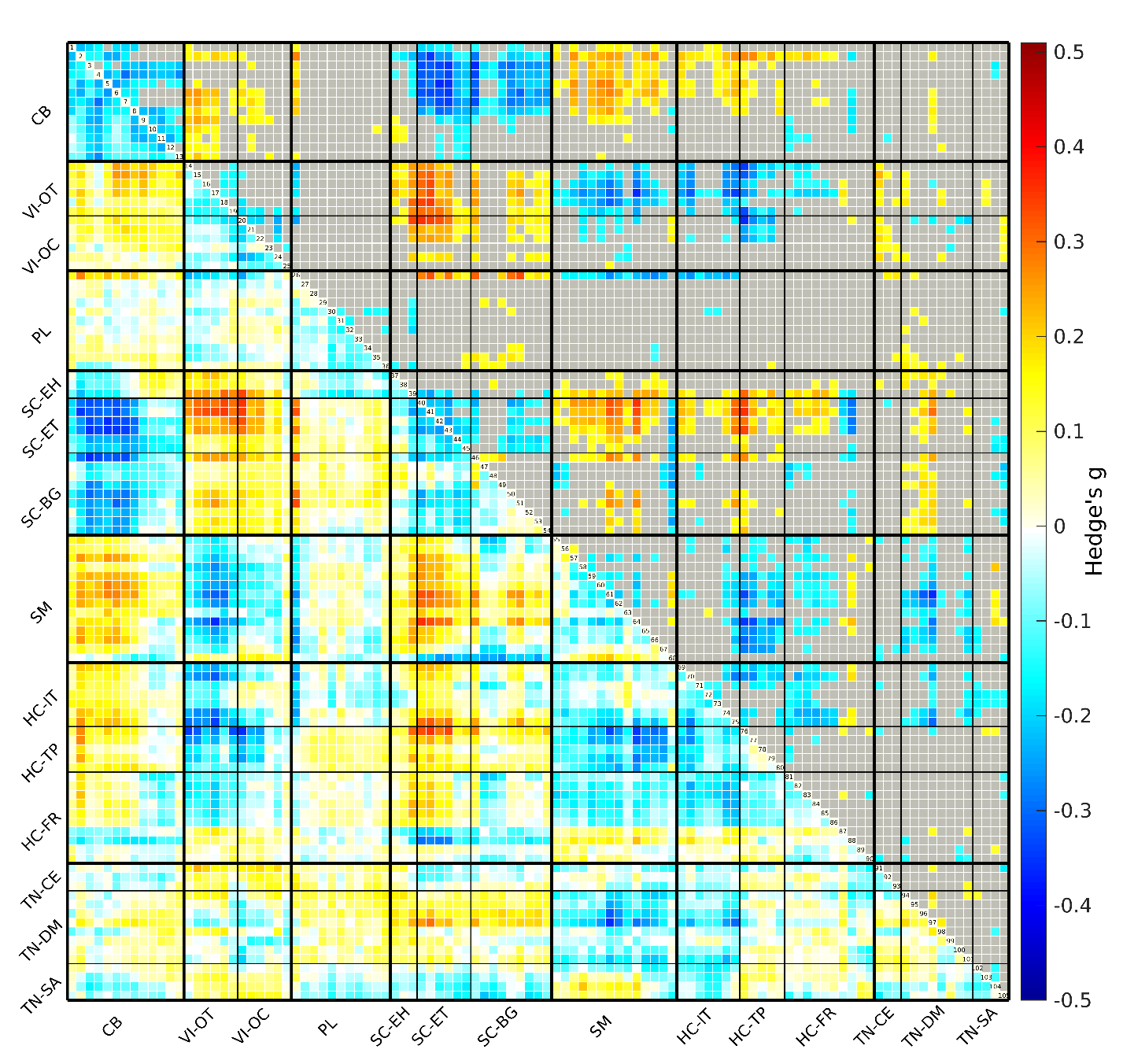

**Figure S3** | **Schizophrenia-Control FNC Group Differences Controlling for CPZ.** Group comparisons controlling for CPZ for each combination of the 105 NeuroMark intrinsic connectivity networks (ICNs) is displayed in the matrix. The upper triangle displays only significant (FDR corrected *q* < .05) case-control group differences in functional network connectivity (FNC) with non-significant pairs grayed out. The heatmap uses red to represent an increase in FNC in schizophrenia relative to controls and blue is used to represent a relative decrease in schizophrenia. ICNs are grouped by their domain-subdomain labels, consistent with the NeuroMark_fMRI_2.2 template: cerebellar (CB), visual-occipitotemporal (VI-OT), visual-occipital (VI-OC), paralimbic (PL), subcortical-extended hippocampal (SC-EH), subcortical-extended thalamic (SC-ET), subcortical-basal ganglia (SC-BG), sensorimotor (SM), higher cognition-insular-temporal (HC-IT), higher cognition-temporoparietal (HC-TP), higher cognition-frontal (HC-FR), triple network-central executive (TN-CE), triple network-default mode (TN-DM), and triple network-salience (TN-SA).

Supplementary Information S8. PANSS Total

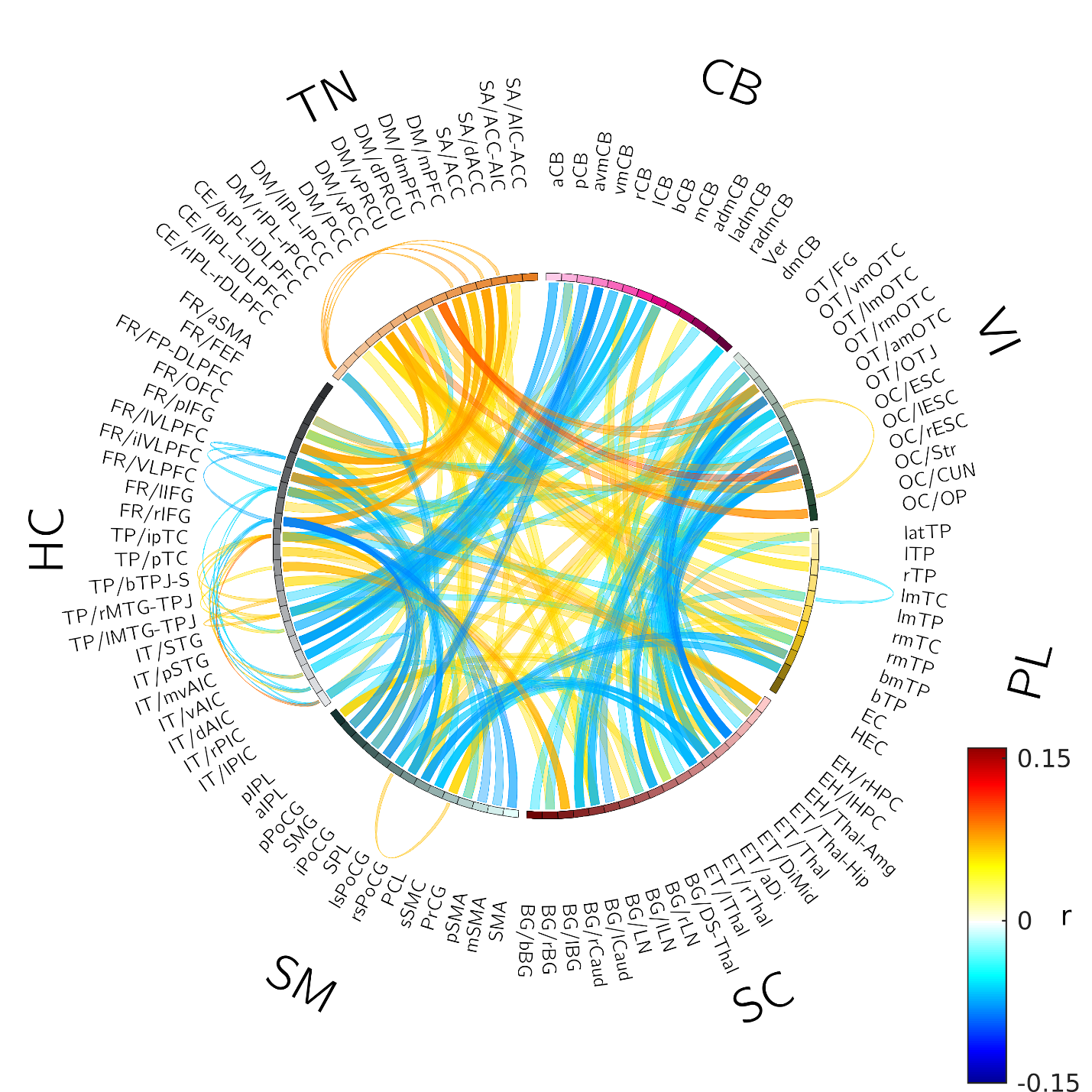

**Figure S4** | **FNC and PANSS Total**. Each of the functional network connectivity (FNC) pairs with significant (FDR corrected *q* < .05) group differences between schizophrenia and control groups in the main analysis were tested for correlations with PANSS total. The connectogram displays significant (*p* < .05) FNC-PANSS total correlations calculated for a reduced subset of individuals with PANSS total scores (N = 1222). In the heatmap, red represents a positive correlation, and blue represents a negative correlation. ICNs are grouped by their domain-subdomain labels, consistent with the NeuroMark_fMRI_2.2 template: cerebellar (CB), visual-occipitotemporal (VI-OT), visual-occipital (VI-OC), paralimbic (PL), subcortical-extended hippocampal (SC-EH), subcortical-extended thalamic (SC-ET), subcortical-basal ganglia (SC-BG), sensorimotor (SM), higher cognition-insular-temporal (HC-IT), higher cognition-temporoparietal (HC-TP), higher cognition-frontal (HC-FR), triple network-central executive (TN-CE), triple network-default mode (TN-DM), and triple network-salience (TN-SA).

Supplementary Information S9. Sample Size Effects Analysis

**Table S15** | **Sample Size Effects Analysis.** In order to further investigate the effects of reduced sample size on each QC analysis we repeated each analysis 1,000 times, randomly sampling subsets of equivalent sample size without replacement from the main subset for each analysis. For example, from the 562 subjects with CPZ scores, 259 subjects were retained in the QC analysis, therefore, 259 subjects were randomly sampled without replacement from the 562 subjects with CPZ scores and the analysis was repeated on this random subset. Each analysis was repeated with 1,000 random subsets and the results were averaged across iterations. Specifically, for each variable, the number of subjects in the main subset is reported, then the number of subjects in both the QC and random subset analyses, then the percentage of significant FNC pairs from the main analysis which replicated in the QC subset, then the average percentage of significant FNC pairs from the main analysis which replicated across 1,000 random subsets, then the Pearson correlation coefficient *r* between the results of the main and QC analysis, and the average Pearson correlation coefficient *r* between the results of the main and random subsets are presented.

| **Variable** | **Main Subset N** | **QC/Random Subset N** | **% FNCs Significant in QC Subset** | **% FNCs Significant in Random Subsets** | ***r* in QC Subset** | **Avg *r* across Random Subsets** |
| --- | --- | --- | --- | --- | --- | --- |
| CPZ | 562 | 259 | 34.93% | 31.77% | 0.728 | 0.716 |
| PANSS_pos | 1091 | 630 | 47.20% | 37.95% | 0.748 | 0.764 |
| PANSS_neg | 1091 | 630 | 22.41% | 41.46% | 0.645 | 0.785 |
| PANSS_gen | 1090 | 629 | 29.86% | 37.76% | 0.761 | 0.783 |
| PANSS_total | 1222 | 715 | 31.56% | 38.93% | 0.770 | 0.787 |
| WM_z | 2328 | 1533 | 33.09% | 47.24% | 0.690 | 0.804 |
| ProcSpd_z | 1753 | 1021 | 47.47% | 41.57% | 0.809 | 0.804 |
| VerbL_z | 1543 | 868 | 38.24% | 45.05% | 0.745 | 0.798 |
| RProbS_z | 1543 | 868 | 31.58% | 39.05% | 0.746 | 0.772 |
| VisL_z | 443 | 175 | 53.42% | 27.39% | 0.759 | 0.700 |
| AttVig_z | 441 | 176 | 15.04% | 26.75% | 0.470 | 0.661 |
| SoCog_z | 155 | 56 | 28.45% | 24.45% | 0.503 | 0.569 |
| dur_ill_yrs | 792 | 416 | 36.36% | 40.64% | 0.747 | 0.819 |
| AO | 792 | 416 | 36.19% | 40.20% | 0.657 | 0.719 |
| **Group** | 2656 | 1727 | 86.68% | 87.90% | 0.970 | 0.970 |

Supplementary Information S10. NeuroMark-SZ Template

To facilitate future efforts, we have released the schizophrenia-control group difference effect sizes of our analysis of rsfMRI for 2,656 individuals (1,248 with schizophrenia) as the NeuroMark-SZ template. This includes a detailed (105 ICN by 105 ICN) effect size matrix corresponding with the NeuroMark 2.2 ordering ('NeuroMark_SZ_effect_size_matrix.csv'), as well as a binary significance matrix indicating the FNC features with significant group differences (FDR q < .05) with ‘1’ and non-significant features with ‘0’ ('NeuroMark_SZ_FDR_sig_matrix.csv'). This resource is publicly available for download at <https://github.com/trendscenter/gift/blob/master/doc/web/wiki/neuromark/functionalMRI/Neuromark_Template_2.2_fMRI_SZ.md>.
